# Engineered extracellular vesicle decoy receptor-mediated modulation of the IL6 trans-signalling pathway in muscle

**DOI:** 10.1101/2020.06.09.142216

**Authors:** Mariana Conceição, Laura Forcina, Oscar P. B. Wiklander, Dhanu Gupta, Joel Z. Nordin, Graham McClorey, Imre Mäger, André Görgens, Per Lundin, Antonio Musarò, Matthew J. A. Wood, Samir EL Andaloussi, Thomas C. Roberts

## Abstract

The cytokine interleukin 6 (IL6) is a key mediator of inflammation that contributes to skeletal muscle pathophysiology. IL6 activates target cells by two different mechanisms, the classical and transsignalling pathways. While classical signalling is associated with the anti-inflammatory activities of the cytokine, the IL6 trans-signalling pathway mediates chronic inflammation and is therefore a target for therapeutic intervention. Extracellular vesicles (EVs) are natural, lipid-bound nanoparticles, with potential as targeted delivery vehicles for therapeutic macromolecules. Here, we engineered EVs to express IL6 signal transducer (IL6ST) decoy receptors to selectively inhibit the IL6 trans-signalling pathway. The potency of the IL6ST decoy receptor EVs was optimized by inclusion of a GCN4 dimerization domain and a peptide sequence derived from syntenin-1 which targets the decoy receptor to EVs. The resulting engineered EVs were able to efficiently inhibit activation of the IL6 transsignalling pathway in reporter cells, while having no effect on the IL6 classical signalling. IL6ST decoy receptor EVs, were also capable of blocking the IL6 trans-signalling pathway in C2C12 myoblasts and myotubes, thereby inhibiting the phosphorylation of STAT3 and partially reversing the anti-differentiation effects observed when treating cells with IL6/IL6R complexes. Treatment of a Duchenne muscular dystrophy mouse model with IL6ST decoy receptor EVs resulted in a reduction in STAT3 phosphorylation in the quadriceps and gastrocnemius muscles of these mice, thereby demonstrating *in vivo* activity of the decoy receptor EVs as a potential therapy. Taken together, this study reveals the IL6 trans-signalling pathway as a promising therapeutic target in DMD, and demonstrates the therapeutic potential of IL6ST decoy receptor EVs.

## Introduction

Extracellular vesicles (EVs), such as exosomes and microvesicles, are endogenous nano-sized vesicles that are released by many cell types and mediate intercellular communication under both physiological and pathophysiological conditions [1]. EVs have been extensively studied as delivery vehicles for therapeutics, as they can transport a multitude of molecules between cells (including nucleic acids, proteins, and lipids) [2–6]. Aside from their ability to carry bioactive molecules, EVs derived from specific cell types such as mesenchymal stem cells have been successfully used for clinical applications, as they exhibit other potentially beneficial features (such as low immunogenicity, inherent anti-inflammatory, and pro-regenerative properties [7–12]). It was previously shown that EVs can be engineered to carry biologically active molecules (both targeting ligands and/or therapeutic proteins), by fusing them to EV-associated proteins [3, 13–16]. This strategy can be used to express decoy receptors on the surface of EVs, with the capability of binding to specific signalling molecules with high affinity and specificity, and thereby blocking their intercellular signalling cascades [16, 17]. This approach presents the opportunity for the modulation of inflammation by targeting key cytokines, such as interleukin 6 (IL6).

IL6 is a multifunctional cytokine, with both pro- and anti-inflammatory properties, that exerts its biological activities primarily through two different mechanisms; the classical- and trans-signalling pathway [18, 19]. (Notably, an additional mode of IL6ST activation was recently identified, the IL6 trans-presentation by dendritic cells [20]). In the case of the classical signalling pathway, associated with the anti-inflammatory activities of the cytokine, IL6 first binds to the membrane-bound interleukin 6 receptor (IL6R). The complex of IL6 and IL6R further associates with IL6ST (IL6 Signal Transducer, also known as gp130), which then dimerizes and initiates signal transduction via the activation of cytoplasmic tyrosine kinases and STAT3 (signal transducer and activator of transcription 3) [19]. Although the expression of the IL6R is restricted to relatively few cell types (e.g. hepatocytes, neutrophils, and macrophages), IL6 was still found to activate cells that lack membrane-bound IL6R [19], an observation which led to the discovery of the trans-signalling pathway. In this alternative pathway, IL6 first binds to a naturally-occurring, soluble-form of the IL6 receptor (sIL6R, primarily generated by alternative splicing or proteolytic processing of the IL6R protein) in the extracellular space, and subsequently the IL6/IL6R complex binds to and activates IL6ST on the surface of cells that do not express membrane-bound IL6R [21]. The IL6 trans-signalling pathway has been linked to the pro-inflammatory activities of the cytokine, and it is believed to mediate chronic inflammation [22–24]. Since IL6ST is ubiquitously expressed, the activity of the IL6 is greatly expanded relative to the classical pathway [25]. Furthermore, soluble forms of IL6ST were found in human body fluids and these were shown to be able to act as natural inhibitors of the IL6 trans-signalling pathway [26].

Inflammation is a characteristic feature of multiple muscle pathologies, including Duchenne muscular dystrophy (DMD) [27, 28]. DMD is caused by mutations which lead to the loss of the dystrophin protein, which is important for maintaining muscle membrane integrity [29]. The lack of this structural protein results in membrane damage, massive infiltration of immune cells, chronic inflammation, impaired regeneration, and severe skeletal muscle degeneration, which are hallmark features of DMD pathology [30–35]. Inflammation is therefore a major target for the treatment of DMD and consequently, almost all DMD patients are treated with corticosteroids (e.g. prednisolone and deflazacort). However, chronic treatment with corticosteroids have demonstrated some limited efficacy in terms of delaying the loss of ambulation [36, 37], and is associated with a number of undesirable side effects such as weight gain, cushingoid features, behavioral problems, osteoporosis and increased risk of infections [37, 38]. As such, experimental therapies (such as vamolorone and edasalonexent) which reduce inflammation without the detrimental side effects of corticosteroids, are in late-stage clinical development [39, 40].

It is expected that blocking specific mediators of the inflammatory response, such as the IL6 transsignalling pathway, might provide another alternative to steroidal drugs, with increased efficacy and decreased side effects. Multiple lines of evidence suggest that targeting the IL6 pathway in muscle-related pathologies might be beneficial; (i) chronic elevation of IL6 promotes skeletal muscle wasting [41–43], (ii) transient inhibition of the STAT3 signalling pathway stimulates muscle regeneration in aged and dystrophic mice [44, 45], (iii) elevated levels of IL6 results in an increase in myostatin in muscle and consequent loss of muscle mass [46], (iv) serum levels of IL6 are increased in DMD patients and dystrophic mouse models [47–49], (v) the use of corticosteroids decreases the levels of IL6 in DMD patients and a mouse model of DMD, suggesting that the benefits of targeting the IL6 pathway are partially responsible for the positive effects of corticosteroids in DMD [48, 50], (vi) targeting the IL6 pathway using IL6R neutralizing antibodies attenuates muscular dystrophy in mouse models of DMD [48, 51] and (vii) overexpression of IL6 in a DMD mouse model exacerbates the dystrophic muscle phenotype and can more faithfully recapitulate patients disease [49].

Here, we first explored the relevance of the IL6 trans-signalling pathway in myogenic cultures and mouse models of DMD, which led to the identification of this pathway as the dominant IL6 signalling pathway in muscle cells. We then engineered EVs to express signalling incompetent IL6ST decoy receptors, which could specifically block the IL6 trans-signalling pathway. This study demonstrates the activity of IL6ST decoy receptor EVs in reporter cell lines, muscle cell cultures, and *in vivo* muscle.

## Materials and Methods

### Cell culture

C2C12, RAW264.7, HEK293T (all from American Type Culture Collection, Manassas, VA, USA), immortalized human bone marrow derived mesenchymal stromal cells (MSCs) [52], HEK-Blue IL6 (InvivoGen, San Diego, CA, USA), and STAT3 Luciferase HeLa cells (Signosis, Santa Clara, CA, USA) were grown at 37°C in 5% CO_2_. C2C12 cells were maintained in growth media (GM) Dulbecco’s Modified Eagle’s Media supplemented with 15% fetal bovine serum (FBS) and 1× antibiotic-antimycotic (Ab-Am, all Life Technologies, Waltham, MA, USA). RAW264.7 and HEK293T were maintained in GM consisting of DMEM supplemented with 10% FBS and 1× Ab-Am. MSCs were cultured in GM made of low-glucose Roswell Park Memorial Institute (RPMI-1640) supplemented with GlutaMAX and HEPES (both Life Technologies), 10% FBS and 1× Ab-Am. HEK-Blue IL6 and STAT3 Luciferase HeLa reporter cell lines were cultured and used following the manufacturer’s instructions. All cell cultures were confirmed to be negative for mycoplasma contamination in weekly testing.

### STAT3 signalling induction in reporter cell lines

For HEK-Blue IL6 experiments, a cell suspension at a density of 560,000 cells/ml was prepared in GM. A mix of recombinant 10 ng/ml IL6 or IL6/IL6R complexes (R&D Systems, Minneapolis, MN, USA), with or without EVs, was prepared in 10 μl of phosphate-buffered saline (PBS) and added to the corresponding well of a 96 well-plate. Next, 90 μl of the cell suspension was added per well, and cells were incubated at 37°C in 5% CO_2_ for 24 hours. After this time, 20 μl of supernatant of secreted embryonic alkaline phosphatase (SEAP) expressing cells or unconditioned media (negative control), was transferred to a flat bottom 96-well plate. 180 μl of resuspended Quanti Blue (InvivoGen) was added to each well and mixed thoroughly. Plates were incubated at 37°C for 2-3 hours and SEAP levels were determined by absorbance at 620-655 nm.

For HeLa STAT3 Luciferase experiments, cells were seeded in GM at a density of 30,000 cells per well in a 96 well-plate. The next day, cells were treated for 7 hours with IL6/IL6R complexes (10 ng/ml), in the presence or absence of EVs. Media was then removed, cells were washed gently with PBS, and incubated with Glo lysis buffer (Promega, Madison, WI, USA) for 5 minutes at room temperature. 50 μl were transferred to a white opaque 96 well plate, and 50 μl of Bright-Glo luciferase substrate (Promega) was added to each well.

A Victor 3 Multilabel Counter Model 1420 (Perkin Elmer, Waltham, MA, USA) was used for both luminescence and spectrophotometry measurements.

### C2C12 treatments

C2C12s myoblasts were seeded in GM and switched to differentiation media (DM, DMEM supplemented with 2% horse serum) for 5-7 days in order to induce differentiation into myotubes. Myotubes were then treated for 24 hours with recombinant IL6 or IL6/IL6R complexes (at 1 or 2.5 nM, as appropriate) and cells were collected for western blot (WB) or reverse transcriptase quantitative PCR (RT-qPCR). C2C12 myoblasts or myotubes were treated with IL6/IL6R complexes (30 ng/ml) for 6 hours in the presence or absence of WT-EVs or IL6ST-EVs (1×10^10^ particles), and then collected for WB to quantify STAT3 phosphorylation. For immunofluorescence experiments, C2C12 cells were seeded in GM on day 0 and treatments were started on day 1 in GM. On day 2, media was changed to DM and cells were cultured for further 72 hours in the presence of treatment. STAT3 was induced by treatment with IL6/IL6R complexes (30 ng/ml), and uninduced cells were used as a control.

### Animal experimentation

Experiments involving wild-type (WT, either C57/BL10 for experiments comparing DMD mouse models or C57/BL6 for biodistribution experiments), *mdx,* or dystrophin/utrophin double-knockout (DKO) mice were carried out at the Biomedical Sciences Unit, University of Oxford, in accordance with procedures approved by the UK home office (project licence 30/2907). *mdx* mice over-expressing IL6 (*mdx*/IL6) were generated as described previously [49]. Experiments using *mdx*/IL6 mice were performed at Sapienza University (Rome), and were approved by the ethics committee of Sapienza University of Rome and according to the Italian Law on the Protection of Animals.

Mice were sacrificed by escalating CO_2_ concentration, muscles were macrodissected, and tissues stored at −80°C for further processing.

For biodistribution experiments, purified MSC IL6ST-EVs were incubated with 50 μM DiR lipophilic tracer (1,1-dioctadecyl-3,3,3,3-tetramethylindotricarbocyanine iodide, D12731, Invitrogen) with gentle agitation, at 4°C overnight. Free DiR was then separated from EV-bound DiR, by size exclusion liquid chromatography as described below. WT (11 weeks-old) or *mdx* (6 weeks-old) mice were treated with a single subcutaneous injection of DiR-labelled IL6ST-EVs (8×10^9^ EVs/gram). Animals were sacrificed 24 hours after administration and organs harvested to visualize fluorescence *ex vivo.* Images were acquired using an IVIS Lumina II instrument and analysed using Living Image software version 3.2 (both Perkin Elmer).

### Satellite cell isolation

Mouse satellite cells were isolated by enzymatic dissociation of skeletal muscles as described previously [53]. A pre-plating technique was used, that takes advantage of the fact that fibroblasts adhere to the plastic much more readily than do myoblasts. Briefly, mouse hind limb muscles (tibialis anterior, quadriceps and gastrocnemius) were minced and incubated in a 1 mg/ml collagenase/dispase solution (Roche, Basel, Switzerland) with agitation at 37°C for 1 hour. An equal volume of plating media (DMEM GlutaMAX with 20% FBS, 1× Ab-Am, and 2% chicken embryo extract from P.A.A. Laboratories, Yeovil, UK) was added and the homogenate was sequentially filtered through a 100 μm cell strainer, followed by a 40 μm cell strainer (both BD Falcon, Franklin Lakes, NJ, USA). The cell suspension flow-through was pre-plated in a 10 cm dish and incubated for 1 hour at 37°C in 5% CO_2_, to allow the fibroblasts to adhere. The media was then carefully removed from the 10 cm dish and the enriched myoblasts population was seeded in a 6 well-plate coated with Matrigel (Sigma-Aldrich, St. Louis, MO, USA) in plating media. Cells were collected 2 days later for analysis.

### Plasmid constructs and cloning

The mouse interleukin 6 signalling transducer (IL6ST) and tumor necrosis factor receptor (TNFR) receptor decoy designs were synthesized by Gen9 Inc. (Cambridge, MA, USA) and cloned into a pLEX vector backbone [54]. Both constructs were then subcloned into the pHAGE lentiviral overexpression plasmid, containing a puromycin-resistance gene for selection. The lentiviral transfer vector plasmid, pSPAX2 packaging plasmid, and pVSV-G envelope plasmid were obtained from Addgene (Watertown, MA, USA).

### Production of lentiviral vectors and stable cell lines

Second generation lentiviral vectors encoding for the various receptor decoy constructs were produced by transient co-transfection of HEK293T cells using a three-plasmid system. Briefly, HEK293T cells were transfected with polyethyleneimine (PEI) mixed with the lentiviral transfer vector plasmid, packaging plasmid, and envelope plasmid in OptiMEM (Life Technologies) for 5 to 6 hours (PEI:total pDNA μg ratio 2:1). After this time, media was changed to GM and the lentiviral vectors were harvested 2 days after transfection by ultracentrifugation of conditioned media (70,000 *g* for 2 hours). The pelleted lentiviral particles were then resuspended in PBS containing 0.5% bovine serum albumin (BSA).

To obtain stable cell lines for the different constructs, HEK293T cells or MSCs were transduced with 2 or 4 μl, respectively, of the concentrated lentiviral vector stock in the presence of polybrene (Sigma-Aldrich). To achieve stable expression of the vectors, infected cells were selected with puromycin, starting at 48 hours after infection, for at least 10 days before performing experiments.

### EV production, purification, and characterization

To isolate EVs, cells were seeded in 150×20 mm dishes in GM until confluent, for the majority of the experiments. After this time, cells were washed with PBS and media was changed to OptiMEM for 48 hours.

For *in vivo* experiments, FiberCell hollow fiber bioreactors with 20 kDa cutoff hydrophilic fibers (KDBIO, Berstett, France) were used for large-scale production of MSC-derived EVs, according to manufacturer’s instructions. Briefly, ~5×10^8^ MSCs (WT or stably-expressing the 2^nd^ generation IL6ST decoy receptor) were seeded in the extra-capillary space (ECS) of the bioreactor using serum-free media. Glucose consumption from the intra-capillary space (ICS) media was monitored every day, and the complete GM detailed above was changed when glucose levels were depleted by 50%. EV harvests started one week after seeding of the cells, by gently flushing the ECS with 25 ml of serum-free RPMI GlutaMAX media supplemented with Ab-Am. Two to three extra harvests were performed in which media from the ICS was withdrawn and flushed back and forth between the two syringes to collect additional EVs. Harvests were performed every 2 or 3 days, up to two months after seeding the cells.

The protocol used to isolate EVs was based on the ultrafiltration combined with size-exclusion liquid chromatography method (UF-SEC-LC) that we have described previously [55]. Conditioned media (CM) from plates or bioreactors was first spun at 700 *g* for 5 minutes to remove cells, and then at 4,000 *g* for 15 minutes to remove cell debris. The resulting supernatant was then passed through a 0.22 μm filter, where possible, to eliminate larger vesicles. When this was not possible, a 10,000 *g* centrifugation for 30 minutes after the concentration step was performed instead. The resulting CM was then concentrated to approximately 10 ml using the Vivaflow 50R tangential flow device (Sartorius, Göttingen, Germany) with a 100 kDa cut-off membrane. CM was further concentrated to approximately 2 ml using 100 kDa cut-off Amicon spin filters (Merk Millipore, Burlington, MA, USA). Concentrated samples were then loaded onto a Sepharose 4 Fast Flow gel filtration column, connected to a ÄKTA Pure apparatus equipped with a UV 280 nm flow cell (all from GE Healthcare, Chicago, IL, USA). EV fractions were collected based on UV absorbance, pooled and concentrated using 30 kDa cut-off Amicon spin filters, when necessary.

The resulting EVs were characterized by nanoparticle tracking analysis (NTA), using the Nanosight NS500 instrument (Malvern, Worcestershire Beacon, UK). To measure the size distribution and EV concentration, quantifications were performed with particle counts in the range of 5×10^8^ to 1×10^9^ EVs per ml. For all our recordings, a camera level setting of 12-14 was used, and three 30-second videos were recorded for each sample, with a delay of 7 seconds between each recording. All particle tracking analyses were performed in duplicate.

To visualize EVs using transmission electron microscopy (TEM), 10 μl of EV sample was applied to freshly glow discharged carbon formvar 300 mesh copper grids (Agar Scientific, London, UK) for 2 minutes. The grid was then blotted dry with filter paper and stained with 2% uranyl acetate for 10 seconds, then blotted and air dried. Grids were imaged using a FEI Tecnai 12 TEM at 120 kV using a Gatan OneView CMOS camera (Gatan, Pleasanton, CA, USA).

### Flow cytometry

C2C12 and RAW264.7 cells were collected, washed with PBS and antibody labelling was performed in live cells using PBS supplemented with 2% FBS (PBS/2%FBS). Details of the antibodies are listed in **Table S1**. Cells were washed twice with PBS after labelling and resuspended in a final volume of 500 μl of PBS/2% FBS for flow cytometry analysis. Expression of cell surface receptors was determined using a FACS Calibur instrument (BD Biosciences, San Jose, CA, USA). A total of 10,000 events per sample were acquired. Data were analysed using the Flowing Software application (Turku University, Finland).

### Western blot

C2C12 cells were harvested in RIPA lysis buffer (Life Technologies) in the presence of protease and phosphatase inhibitor cocktails (Roche). The lysate was incubated on ice for 1 hour and then centrifuged to remove cell debris. To analyse muscle samples, a small piece from each muscle was homogenized in lysis buffer (1 mM EDTA, 5 mM sodium fluoride, 0.5% Triton, and 6 M urea) in the presence of protease and phosphatase inhibitor cocktails (Roche). Mechanical homogenization of the tissue was performed using a Precellys 24 tissue homogenizer (Bertin Instruments, Bretonneux, France). Protein concentration was determined by BCA assay (Thermo Fisher Scientific, Waltham, MA, USA). Fifty micrograms of total denatured protein were then separated by SDS-PAGE using precast NuPAGE 4-12% Bis-Tris protein gels (Invitrogen, Waltham, MA, USA) followed by electrotransfer onto polyvinylidene difluoride (PVDF) membranes. Membranes were blocked in 5% non-fat milk and immunoblotting performed using the antibodies listed in **Table S1**. Chemiluminescence or fluorescence signal was measured, as appropriate, using the Odyssey Fc Imaging System (LI-COR Biosciences, Lincoln, NE, USA). Quantitative analysis was carried out using Image Studio Lite software version 5.2 (LI-COR Biosciences). GAPDH or vinculin were used as reference proteins to control for equal protein loading.

### Transcript analysis

Total RNA from C2C12 and satellite cells was extracted using the RNeasy Mini Kit (Qiagen, Hilden, Germany) according to manufacturer’s instructions. To isolate total RNA from muscle, TRIzol (Life Technologies) was used followed by column purification. Briefly, a small piece from each muscle was mechanically homogenized in TRIzol, using the Precellys homogenizer. The mixture was then transferred to a new tube, and chloroform was added. Samples were centrifuged at 11,000 *g* (15 minutes, at 4°C) and the aqueous phase was transferred to a new tube. RNA was precipitated by adding one volume of 70% ethanol and the purification proceeded following the RNeasy Mini Kit protocol. DNase treatments were performed in-column for all samples.

For most experiments, one microgram of total RNA was reverse transcribed using the High Capacity cDNA Reverse Transcription Kit (Applied Biosystems, Foster City, CA, USA), according to the manufacturer’s protocol (satellite cells experiments used 200 ng of total RNA). RT-qPCR was performed with FAST SYBR Green Master Mix in a Step One Plus Real-Time PCR System (both Applied Biosystems). Primer sequences (IDT, Coralville, IA, USA) are listed in **Table S2**. Gene-of-interest expression was normalized to expression of a stable reference gene; *Hprt.* Relative mRNA quantification was performed using the Pfaffl method [56], and fold changes reported relative to control samples.

Absolute quantification of *Il6r, Il6st,* or *Hprt* was performed by comparison of sample levels to synthetic DNA standard curves of known copy numbers (IDT, **Table S2**) analysed in parallel.

To detect alternative splicing of the IL6 receptor, mRNA transcripts were amplified via semi-quantitative RT-PCR using the primers listed in **Table S3**. PCR products were then separated by agarose gel electrophoresis, to detect the presence of bands corresponding to the various splice products. RNA from human lymphoblasts and myoblasts, used as a positive control for alternative splicing of the *IL6R,* were kind gifts from Dr. Lara Cravo and Prof. Carlo Rinaldi, respectively.

### Immunofluorescence

To perform myosin heavy chain (MHC) staining, C2C12 cells were fixed with 4% paraformaldehyde, permeabilized with 0.25% Triton X-100, and blocked with 5% bovine serum albumin (BSA), as previously described [57]. Incubations with primary and secondary antibodies were performed in 5% BSA blocking solution (details of antibodies are shown in **Table S1**). Nuclei were stained with Hoechst 33258 (H3569 from Life Technologies, 1:5,000 dilution). Microscopy images were acquired using an EVOS FL microscope (Life Technologies) with a 10× objective lens. Quantification of the MHC positive area was performed using ImageJ (NIH, Bethesda, MD, USA). All fluorescence parameters were kept constant to enable comparisons across images. Myogenic index was defined as the percentage of nuclei within MHC-positive cells. Fusion index was defined as the percentage of nuclei within MHC-positive myotubes containing at least 3 nuclei.

### Statistical analysis

The number of independent experiments is indicated in each figure legend. Comparisons between two groups were tested using an unpaired Student’s *t*-test. Multiple groups were compared using one-way ANOVA combined with Bonferroni’s post *hoc* test. Statistically significant differences are presented at probability levels of \**P*<0.05, \*\**P*<0.01 and \*\*\**P*<0.001. Statistical analyses were performed using GraphPad Prism version 8 (GraphPad Software, Inc., California Corporation, San Diego, CA, USA).

## Results

### The IL6 trans-signalling pathway as a therapeutic target for muscle pathology

Cellular responsiveness towards IL6 or IL6/IL6R complexes is dependent on the ratio between IL6R and IL6ST protein levels in cells [58]. We therefore first evaluated the cell surface expression of these receptors in C2C12 murine skeletal muscle cells (both myoblasts and differentiated myotubes). Flow cytometry data showed that 99.6% of the myoblasts expressed IL6ST while only 6.4% were doublepositive for both IL6R and IL6ST (**Fig. 1A**). For myotubes, approximately 86.2% expressed IL6ST, with only 3.1% expressing both IL6R and IL6ST (**Fig. 1B**). In contrast, 82.6% of RAW264.7 macrophages (which served as a positive control for IL6R expression) were positive for IL6R, while 21.9% expressed both IL6ST and IL6R (**Supplementary Fig. 1A**), indicative of successful detection of IL6R. Isotype controls indicated low levels of non-specific antibody binding (**Supplementary Fig. 1B-D**).

**Figure 1.**
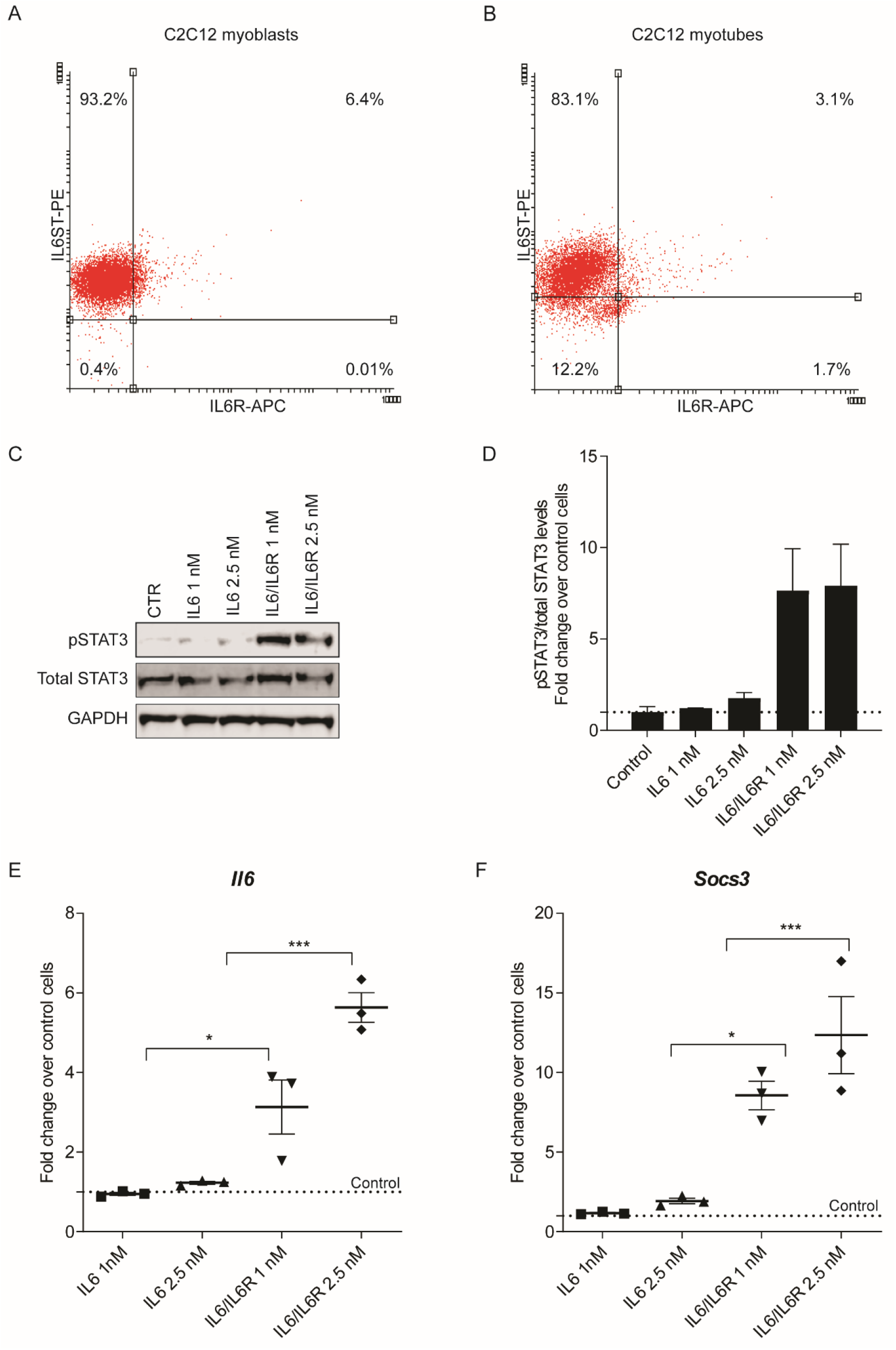
Muscle cells are more responsive to the IL6 trans-signalling pathway than to the classical signalling pathway. C2C12s were incubated with anti-IL6R APC and anti-IL6ST PE, and the expression of IL6R and IL6ST was assessed by flow cytometry. Two-parameter density plots for IL6R and IL6ST expression in C2C12 **(A)** myoblasts and **(B)** myotubes. C2C12 myotubes were treated for 24 hours with recombinant IL6 or IL6/IL6R complexes (1 or 2.5 nM, as indicated), and STAT3 activation was assessed by western blot. **(C)** Representative membrane image for active phospho-STAT3 (pSTAT3, Tyr705), total STAT3, and GAPDH proteins in C2C12 myotubes treated with IL6 or IL6/IL6R complexes. **(D)** Fold change in pSTAT3/total STAT3 ratios in treated cells over untreated (Control) cells. C2C12 myotubes were treated for 24 hours with recombinant IL6 or IL6/IL6R complexes (1 or 2.5 nM, as indicated) and mRNA expression of **(E)** *Il6* or **(F)** *Socs3* determined by RT-qPCR. The dotted line indicates the level of untreated control cells. Gene-of-interest expression was normalized to levels of a stable reference gene; *Hprt.* Values are mean±SEM, *n*=3 independent experiments, \**P*<0.05, \*\*\**P*<0.001 (one-way ANOVA with Bonferroni *post hoc* test).

The activation of the IL6 pathway leads to an increase in STAT3 phosphorylation [59]. Thus, to investigate the differential capability of equivalent concentrations of recombinant IL6 or pre-formed IL6/IL6R complexes to activate the IL6 pathway, the phosphorylation status of STAT3 at tyrosine residue 705 [60] was evaluated in C2C12s myotubes by western blot. Treatment with IL6/IL6R complexes increased STAT3 phosphorylation by ~8-fold, while IL6 alone resulted in only a marginal increase (**Fig. 1C-D**). Similarly, IL6/IL6R complexes activated the expression of IL6-regulated genes much more efficiently than IL6 alone, as shown by the significant increase in the mRNA levels of *Il6* and the suppressor of the cytokine signalling 3 *(Socs3),* evaluated by RT-qPCR (**Fig. 1E-F**). Furthermore, the mRNA levels of other IL6-regulated genes; *Il6r, Il6st, Adam10,* and *Adam17*, were found to be moderately elevated when myotubes were treated with IL6/IL6R complexes, but unchanged in response to IL6 alone (**Supplementary Fig. 2A-D**). ADAM10 and ADAM17 are members of the A Disintegrin And Metalloprotease (ADAM) protein family, that have been implicated in the shedding process that liberates soluble IL6R (sIL6R, an essential effector molecule of the IL6 trans-signalling pathway) from the cell surface. In summary, myogenic cells are responsive to the trans-signalling IL6 pathway, and to a much lesser extent the classical pathway, consistent with the low levels of IL6R observed in these cells.

Skeletal muscle growth and repair is supported by resident stem cells called satellite cells. It was previously reported that high levels of IL6 could induce satellite cell exhaustion, and conversely that transient inhibition of the STAT3 signalling pathway could stimulate muscle regeneration [44, 45]. We therefore aimed to predict the responsiveness of satellite cells to the IL6 classical and trans-signalling pathways, based on the gene expression of *Il6r* and *Il6st.* For this purpose, we isolated satellite cells from WT and *mdx* mice and determined absolute transcript levels for *Il6r, Il6st,* and *Hprt* (reference control gene) by RT-qPCR. Levels of *Il6st* were 23.6 and 29.7 times higher than *Il6r* in WT and *mdx* mice, respectively (**Supplementary Fig. 3**), suggesting that the trans-signalling pathway is the dominant IL6 pathway in muscle satellite cells, similar to the results observed in C2C12 cells.

We next evaluated alterations in the IL6 pathway in various mouse models of DMD; *mdx, mdx* overexpressing human IL6 (*mdx*/IL6), and dystrophin/utrophin double knockout (DKO) mice (utrophin is a dystrophin paralog which can partially compensate for dystrophin absence [61]). The time-points selected to evaluate the IL6 pathway for these various mouse models were based on their respective phenotypes, at ages when inflammation is present. *mdx* mice have been widely used as a preclinical model for DMD. However, the pathology observed in this model is much less severe than that observed in patients, and the lifespan of *mdx* mice is only reduced by ~25% [62]. Additionally, while the first signs of inflammation and muscle necrosis are recognized early in *mdx* mice, inflammation and degeneration/regeneration cycles decline after 12 weeks-of-age [63–65]. In contrast, the DKO mouse is a much more severe model that exhibits greatly reduced survival (~10 weeks) [61, 66]. We therefore investigated alterations in the IL6 pathway at 6.5 weeks-of-age for *mdx* and DKO mice, when inflammation is expected to be present [65], and at 12 weeks-of-age for *mdx*/IL6 mice, as the period of inflammation in this model persists beyond the crisis period observed in the *mdx* mouse [49]. Initially, we analysed diaphragm muscles in all strains, as this muscle was previously reported to exhibit more severe pathology than limb muscles in *mdx* mice [67]. We observed that the mRNA levels of *Il6* were significantly increased in the diaphragm of the three mouse models of DMD, relative to WT animals (**Fig. 2A**). However, the mRNA levels of the IL6 downstream target genes *Socs3, Adam10,* and *Adam17* were only found to be increased in *mdx*/IL6 mice (**Fig. 2B**, data not shown for *mdx* and DKO mice). Additionally, western blot analysis revealed a significant ~4-6 fold (*P*<0.05) increase in phospho-STAT3 in the diaphragm muscles of *mdx,* DKO, and *mdx*/IL6 mice (**Fig. 2C-E**), as well as in hind limb muscles (quadriceps and tibialis anterior, **Supplementary Fig. 4A-C**). However, the levels of sIL6R were only found to be increased in the *mdx*/IL6 mice (~2-3 fold increase, *P*<0.05), for both diaphragm (**Fig. 2F**) and quadriceps (**Supplementary Fig. 4D**).

**Figure 2.**
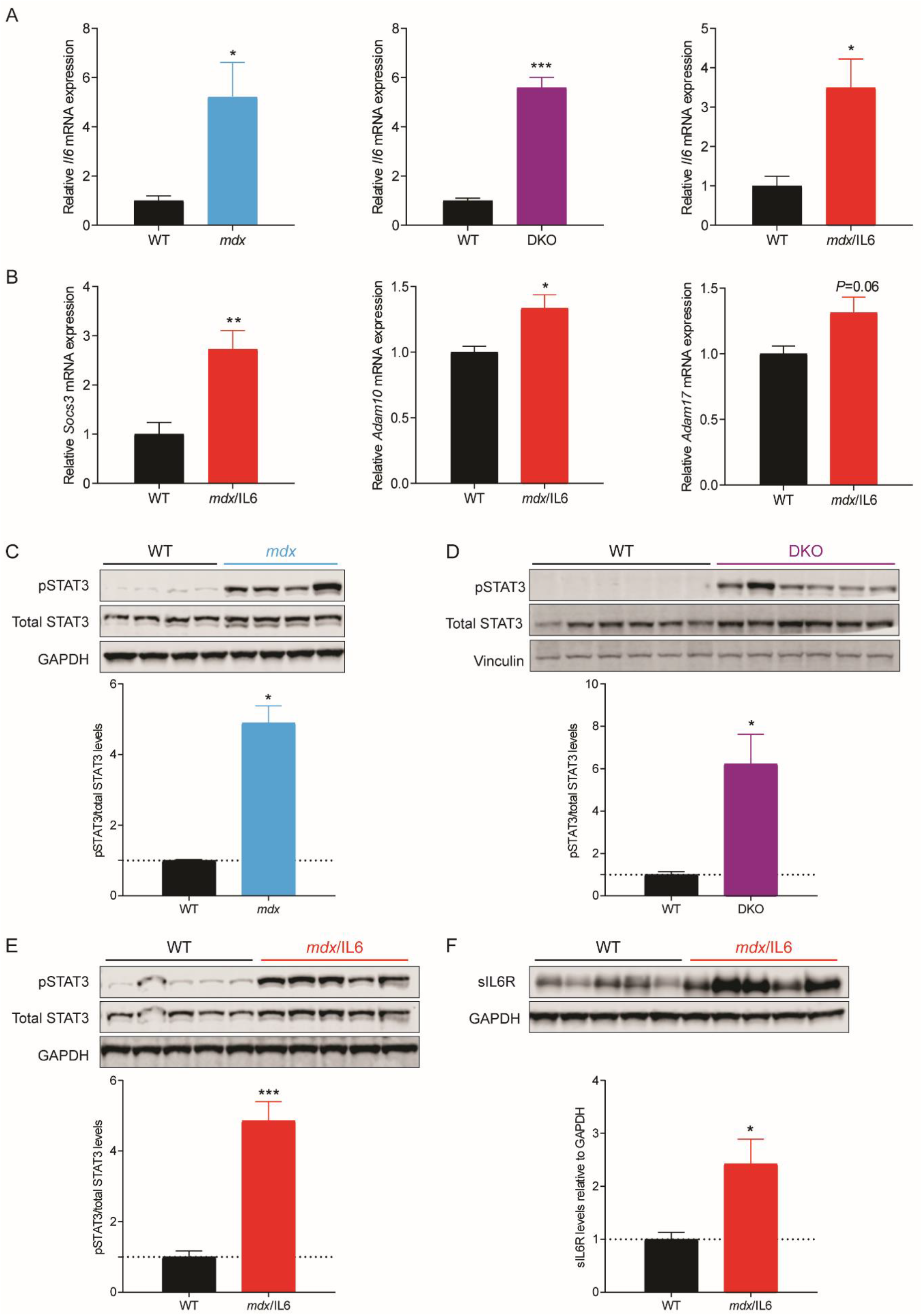
The IL6 pathway is altered in the diaphragm of DMD mouse models. The diaphragm muscle of *mdx,* DKO and *mdx*/IL6 mice was analysed at 6.5 weeks of age for *mdx* and DKO mice, and at 12 weeks of age for *mdx*/IL6 mice and compared with WT (C57BL/10) controls. ****(A)**** *Il6* mRNA levels in the diaphragm of *mdx,* DKO, and *mdx*/IL6 mice as determined by RT-qPCR. **(B)** *Socs3, Adam10,* and *Adam17* mRNA levels in the diaphragm of *mdx*/IL6 mice. For RT-qPCR experiments, the graphs show the fold changes in mRNA levels in the various mouse models of DMD relative to age-matched wild type (WT) mice. Gene-of-interest expression was normalized to levels of a stable reference gene; *Hprt.* Diaphragm muscles from **(C)** *mdx,* **(D)** DKO and **(E)** *mdx*/IL6 mice were assessed by western blot (WB) for active phospho-STAT3 (pSTAT3, Tyr705), total STAT3, and GAPDH or vinculin. **(F)** sIL6R levels in the diaphragm of *mdx*/IL6 mice, assessed by WB. Quantification of WB signal from multiple independent experiments is shown for each analysis. Values are mean+SEM, *n*=5/6 independent experiments for RT-qPCR, or *n*=4-6 independent experiments for WB, \**P*<0.05, \*\**P*<0.01, \*\*\**P*<0.001 (unpaired Student’s *t*-test).

One possible mechanism that can generate sIL6R is alternative splicing, in which the transmembrane domain-encoding region is excluded from the mature *Il6r* mRNA transcript. Here we aimed to evaluate the potential for alternative splicing of the *Il6r* gene in diaphragm muscle lysates from the various DMD mouse models using RT-PCR. Two primer pair combinations were used that span the transmembrane region located in exon 9 [68]. No *Il6r* alternative splice products were observed in any of the mouse models, while alternative splicing (which included previously reported splice isoforms) was observed in human lymphoblasts and myoblasts, that were used as positive controls (**Supplementary Fig. 5**). These findings are consistent with a previous study, which also reported an absence of sIL6R-generating alternative splice isoforms in murine tissues [68].

### Generation and characterization of IL6ST decoy receptor EVs

Having verified the IL6 trans-signalling pathway as a potential disease target in DMD, we next investigated the potential of using engineered EVs to modulate this pathway. Endogenous IL6ST is a 130 kDa membrane protein composed of an N-terminal extracellular domain, a single-pass transmembrane region, and a C-terminal cytoplasmic domain. To generate EVs that could specifically block the IL6 trans-signalling pathway, without affecting the IL6 classical signalling, we designed engineered EVs that express IL6ST decoy receptors at the EV surface, hereafter referred to as IL6ST-EVs. A modified IL6ST transgene was designed in which the majority of the cytoplasmic domain was deleted (40 amino acids of the 277 amino acids were retained), thereby generating a signalling incompetent IL6ST receptor. Two different IL6ST-EV configurations were tested; (i) 1^st^ generation, which displayed the murine signalling incompetent IL6ST receptor, and (ii) 2^nd^ generation, in which the mouse signalling incompetent IL6ST receptor was further fused to a GCN4 leucine zipper domain [69], and to the N-terminal domain of syntenin-1 (SDCBP) (**Fig. 3A**). The leucine zipper domain was included in order to stabilize the receptor complex, improve EV-loading of the modified IL6ST proteins [70], and to promote receptor dimerization, thereby mimicking the functionality of the wild-type receptor on the cell surface. Similarly, fusion with the N-terminal domain of syntenin-1, one of the most frequently detected proteins in EVs that controls the biogenesis of exosomes [71, 72], was included in the design to promote trafficking of the receptors to EVs. The EV-producer cell lines (HEK293T cells and MSCs) were stably transduced with lentiviral vectors encoding the decoy receptors, such that the EVs produced by these cell lines incorporated the respective decoy receptors in their membranes.

**Figure 3.**
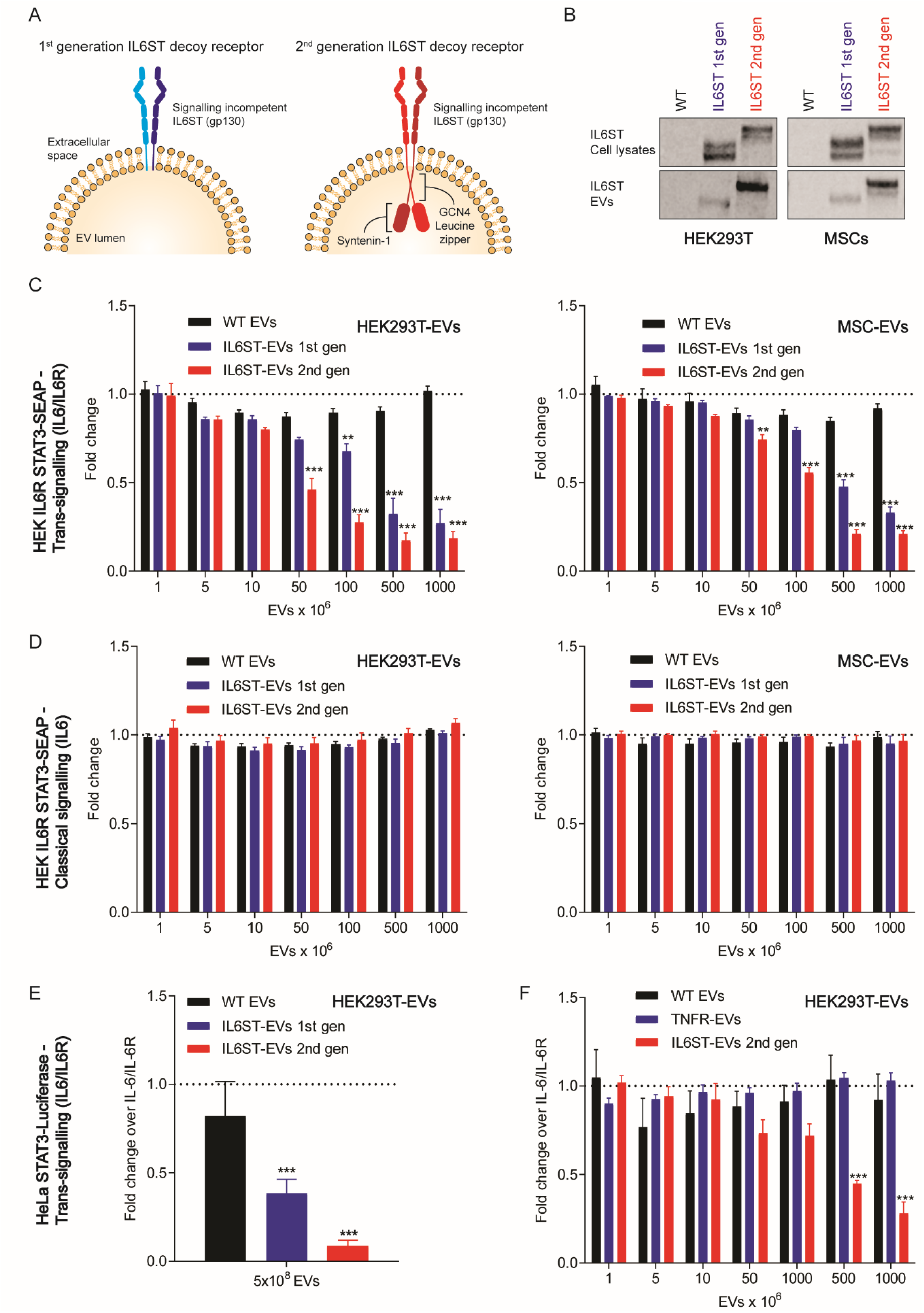
IL6ST-EVs specifically block the IL6 trans-signalling pathway. ****(A)**** Schematic representation of signalling incompetent IL6ST receptor designs in the extracellular vesicle (EV) membrane. 1^st^ generation IL6ST, displays the murine signalling incompetent IL6ST receptor. 2^nd^ generation IL6ST, in which the mouse signalling incompetent IL6ST receptor is fused to a GCN4 leucine zipper domain, and to the N-terminal domain of syntenin-1. **(B)** HEK293T or MSC cultures were transduced with lentiviral vectors encoding for the receptor decoy constructs, and IL6ST expression in cell lysates or isolated EVs was determined by WB. SEAP signal was quantified in HEK IL6R STAT3-SEAP cells induced with 10 ng/mL of either recombinant **(C)** IL6/IL6R complexes or **(D)** IL6, and treated with EVs derived from the corresponding cell line (HEK293T or MSCs) at the indicated doses; WT (used as control), IL6ST 1^st^ generation, or IL6ST 2^nd^ generation. **(E)** Luminescence was determined in HeLa STAT3 Luciferase cells induced with 10 ng/mL of recombinant IL6/IL6R complexes and treated in the presence of 5×10^8^ HEK293T EVs (WT, IL6ST 1^st^ generation, or IL6ST 2^nd^ generation). **(F)** Luciferase signal was measured in HeLa STAT3 Luciferase cells induced with 10 ng/mL of recombinant IL6/IL6R complexes and treated with EVs derived from HEK293T at the indicated doses; TNFR-EVs or WT-EVs (used as a controls), IL6ST 1^st^ generation, or IL6ST 2^nd^ generation. Values are mean+SEM, *n*=3/4 independent experiments, \*\**P*<0.01, \*\*\**P*<0.001 (one-way ANOVA with Bonferroni *post hoc* test).

Vesicles derived from these EV-producer cell lines were isolated by UF-SEC-LC, a scalable method used to obtain pure and intact EVs, devoid of contaminating proteins (**Supplementary Fig. 6A**) [55]. The resulting EVs were characterised using multiple orthogonal methodologies. Nanoparticle tracking analysis showed the presence of particles with a single peak with modal average diameter of 144 nm (**Supplementary Fig. 6B**), and electron microscopy revealed predominantly intact particles with rounded and cup shaped morphology (**Supplementary Fig. 6C**), consistent with the properties of EVs. Furthermore, western blot analysis demonstrated the presence of common EV markers, alix (PDCD6IP), syntenin-1 (SDCBP), and CD81, in EVs, while the endoplasmic reticulum marker protein calnexin (CANX) was enriched in cell lysates (**Supplementary Fig. 6D**).

Immunoblotting for IL6ST demonstrated that the two IL6ST-EV-producer cell lines expressed both IL6ST decoy receptor constructs at similar levels (**Fig. 3B**). Conversely, analysis of the EVs purified from these lines showed that the engineered protein expression was enriched to a greater extent in the 2^nd^ generation IL6ST-EVs, relative to the 1^st^ generation IL6ST-EVs (**Fig. 3B**), likely as a consequence of the inclusion of the additional EV-sorting and receptor dimerization domains. These data demonstrate the successful construction of engineered EV producer cell lines, isolation of modified EVs, and the superior loading of the 2^nd^ generation decoy receptors design.

### IL6ST-EVs selectively block the IL6 trans-signalling pathway

We next sought to determine if IL6ST-EVs could modulate the IL6 trans-signalling in reporter cell lines. HEK Blue IL6R cells (which express secreted embryonic alkaline phosphatase (SEAP) under the regulation of STAT3 responsive elements) were induced with IL6/IL6R complexes, and treated in parallel with either 1^st^ or 2^nd^ generation IL6ST-EVs (isolated from either HEK293T or MSC producer cell lines) and STAT3 inhibitory activity was evaluated by measuring SEAP reporter expression. Treatment with wild-type EVs (WT-EVs) was used as negative control. 1^st^ and 2^nd^ generation IL6ST-EVs, derived from HEK293T or MSC, inhibited IL6/IL6R induced trans-signalling in a dose-dependent manner, while WT-EVs exhibited no effect (**Fig. 3C**). 2^nd^ generation IL6ST-EVs were found to be more potent than the 1^st^ generation decoy-EVs, likely as a result of improved receptor decoy loading on the EV surface and/or due to an increased propensity for receptor dimerization **(Fig. 3C**). We detected up to 83% inhibition in STAT3 activation with 2^nd^ generation IL6ST-EVs (**Fig. 3C**). Importantly, the classical signalling pathway induced by IL6 was not affected by treatments with IL6ST-EVs (**Fig. 3D**). Experiments were repeated in a second reporter cell line (HeLa STAT3-Luciferase cells). While these cells enable facile measurement of STAT3 activation upon treatment of cells with recombinant IL6/IL6R complexes, they have the limitation of not expressing IL6R. As such, HeLa STAT3-Luciferase cells cannot be activated upon treatment with recombinant IL6 alone. Luminescence experiments demonstrated that IL6ST-EVs could inhibit the IL6/IL6R induced trans-signalling pathway, with the 2^nd^ generation IL6ST-EVs performing better (**Fig. 3E**). STAT3 inhibition was dose-dependent, with up to 90% inhibition observed using the 2^nd^ generation IL6ST-EVs (**Fig. 3E-F**). Treatments with control WT-EVs, or EVs engineered to display the tumor necrosis factor receptor (TNFR-EVs) on their surface, had no effect on the induction of the trans-signalling pathway mediated by IL6/IL6R complexes (**Fig. 3F**).

Having demonstrated the efficacy of IL6ST-EVs in reporter cell lines, we next sought to evaluate their activity on IL6 trans-signalling in muscle cells. Given their superior efficacy, only 2^nd^ generation IL6ST-EVs were used in future experiments. C2C12 myoblasts or myotubes were treated with IL6/IL6R complexes in the presence or absence of IL6ST-EVs (or WT-EV controls) and the activation status of STAT3 determined by western blot. IL6ST-EVs inhibited IL6/IL6R complex-mediated STAT3 phosphorylation by ~80% in myoblasts (*P*<0.001) and by at least 50% (*P*<0.01) in myotubes, while WT-EVs had no effect (**Fig. 4A-B**). Three doses of EVs were assessed (1×10^10^, 2×10^10^, and 4×10^10^ EVs per well), although close to maximal effect was observed even with the low dose treatments, and so a clear dose-dependent effect could not be determined from this experiment.

**Figure 4.**
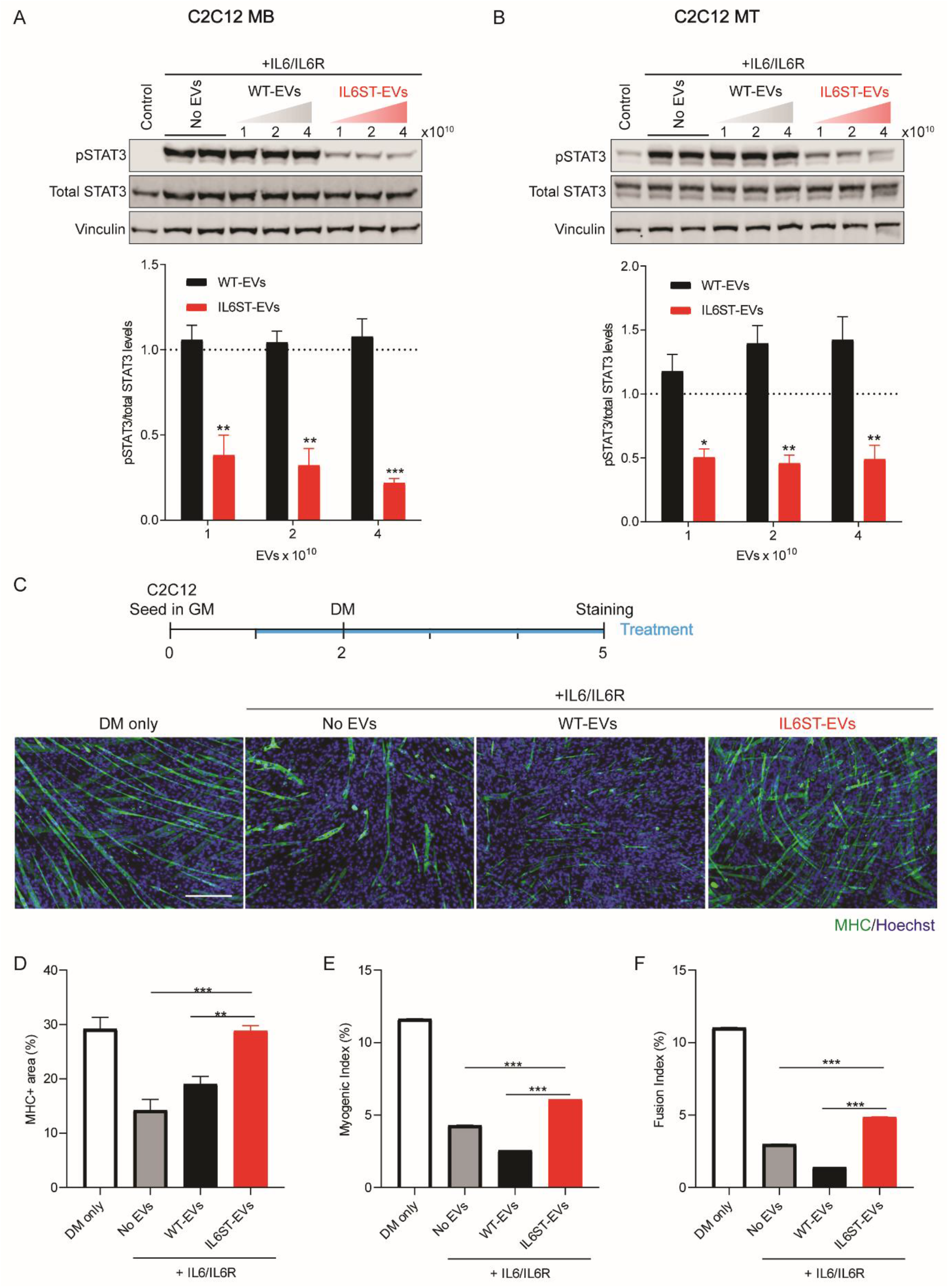
IL6ST-EVs block the IL6 trans-signalling pathway in muscle cells. WB analysis for pSTAT3, total STAT3, and vinculin proteins in **(A)** C2C12 myoblasts (MB) and **(B)** myotubes (MT). C2C12 cultures were induced with recombinant IL6/IL6R complexes (30 ng/ml) and treated with IL6ST-EVs, WT-EVs (concentrations as indicated), or left untreated. Graphs represent the fold change in pSTAT3/total STAT3 ratios for treated cells relative to untreated (control) cells (dotted line). **(C)** C2C12 cells were seeded in growth media (GM) on day 0 and treatments were started on day 1 in GM. On day 2, media was changed to differentiation media (DM) and cells were cultured in the presence of the treatment for further 72 hours. Cells were treated with IL6/IL6R complexes (30 ng/ml), in the presence of absence of 1×10^10^ EVs (WT or IL6ST). Untreated cells (DM only) were used as a control. Myogenic differentiation was assessed by MHC immunofluorescence staining, and quantified by measuring the **(D)** MHC+ area, **(E)** myogenic index, and **(F)** fusion index. Values are presented as mean+SEM of *n*=3 independent experiments for WB, or *n*=4 representative fields of view for microscopy, \**P*<0.05, \*\**P*<0.01, \*\*\**P*<0.001 (one-way ANOVA with Bonferroni *post hoc* test). Images were taken at 10x magnification and the scale bar indicates 200 μm.

To investigate the capacity of IL6ST-EVs to reverse the effects of IL6/IL6R complex-mediated STAT3 activation on myogenic differentiation, C2C12s myoblasts were switched to DM and treated with recombinant IL6/IL6R complexes, in the presence or absence of IL6ST-EVs or WT-EV controls (according to schema shown in **Fig. 4C**). Myogenic differentiation was assessed by immunostaining for myosin heavy chain (MHC), and quantified by calculating the MHC positive area, myogenic index, and fusion index (**Fig. 4C-F**). While treatment of C2C12 cultures with IL6/IL6R complexes decreased myotube formation, concomitant treatment with IL6ST-EVs was able to partially reverse this phenomenon, as observed by MHC immunostaining and quantification of the MHC positive area (**Fig. 4C-D**). Similarly, myogenic- and fusion indices exhibited significant (*P*<0.001) increases towards the level of untreated controls following treatment with IL6ST-EVs, although the level of differentiation did not fully recover to the level of that observed in untreated controls (**Fig. 4E-F**).

Having demonstrated the efficiency of 2^nd^ generation IL6ST-EVs in cell culture, we next sought to test their effectiveness *in vivo.* The biodistribution of IL6ST-EVs was assessed following subcutaneous administration in both WT (C57/BL6) and *mdx* mice. DiR, a lipophilic (membrane intercalating) nearinfrared dye, was used to track the distribution of EVs across multiple murine organs. When comparing EV accumulation in the various muscles, the majority of the signal was observed in the heart, followed by quadriceps and gastrocnemius, while the signal in the diaphragm was low (**Fig. 5A-B**). (Biodistribution results were similar for both WT and *mdx* mice). The evaluation of other organs revealed high levels of IL6ST-EV accumulation in lungs, liver, and kidneys in both WT and *mdx* mice (**Supplementary Fig. 7A-B**). Accumulation in spleen was also observed in WT mice (**Supplementary Fig. 7A**). These results demonstrate that IL6ST-EVs can be successfully delivered to disease-relevant tissues, following subcutaneous administration.

**Figure 5.**
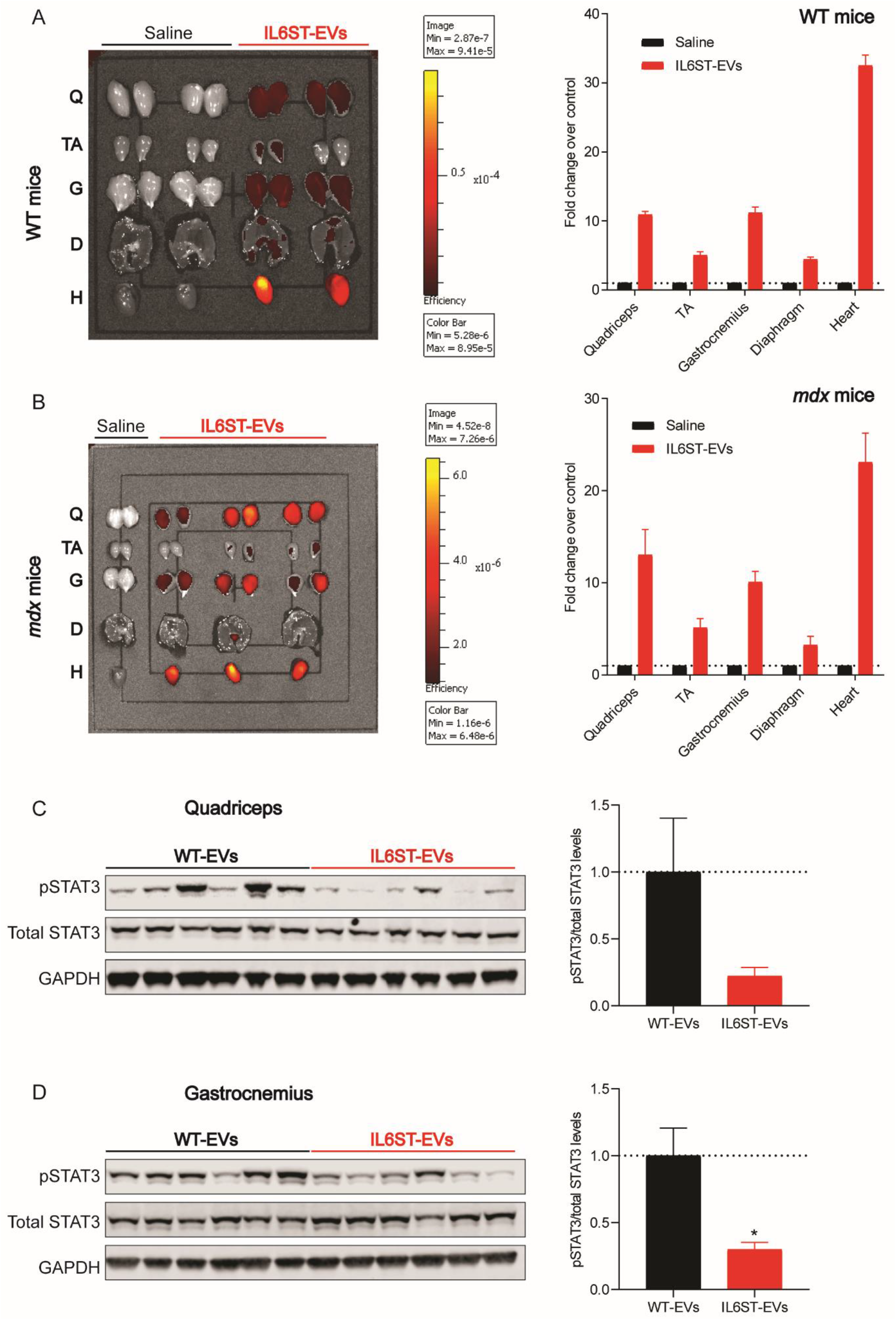
Muscle biodistribution and therapeutic efficacy of IL6ST-EVs. **(A)** WT or **(B)** *mdx* mice were subcutaneously injected with DiR-labelled 2^nd^ generation IL6ST-EVs (8×10^9^ EVs/g) and images were acquired 24 hours after injection using an IVIS Lumina imager. Each image includes quadriceps (Q), tibialis anterior (TA), gastrocnemius (G), diaphragm **(D)** and heart (H) muscles from mice injected with DiR-labelled IL6ST-EVs or saline injected controls. 4 weeks-old *mdx*/IL6 mice were treated with IL6ST-EVs or WT-EVs (1×10^10^ EVs/g) for 2 weeks, 2 injections per week. Mice were sacrificed 3 days after the 4^th^ injection, and hind limb muscles harvested. *mdx*/IL6 treated with IL6ST-EVs or WT-EV controls and protein levels of pSTAT3, total STAT3, and GAPDH determined by western blot analysis in **(C)** quadriceps and **(D)** gastrocnemius muscles. Graphs depict the fold change in pSTAT3/total STAT3 ratios for IL6ST-EV treated animals relative to mice treated with WT-EVs. Values are presented as mean+SEM of *n*=1-3 for biodistribution (as shown in each image), or *n*=5/6 for WB, \**P*<0.05 (unpaired Student’s *t*-test).

To demonstrate *in vivo* activity of the EVs, *mdx*/IL6 mice were treated with 2^nd^ generation IL6ST-EVs or WT-EV controls (two subcutaneous injections of 1×10^10^ EVs/gram/week, for 2 weeks). Animals were sacrificed 3 days after the final (fourth) injection, and the phosphorylation status of STAT3 was evaluated in diaphragm, quadriceps, and gastrocnemius muscles. IL6ST-EVs treatments resulted in a decrease in STAT3 phosphorylation in the quadriceps and gastrocnemius (**Fig. 5C-D**), indicative of successful inhibition of the IL6 signalling pathway. However, no activity was observed in the diaphragm (**Supplementary Fig. 8**), consistent with the biodistribution pattern described above. In summary, these findings demonstrate the potential utility of decoy receptor EVs for the therapeutic modulation of the IL6 trans-signalling pathway *in vivo.*

## Discussion

Chronic inflammation, in which there is prolonged release of pro-inflammatory cytokines, such as IL6, plays a critical role in the pathogenesis of many muscle pathologies and contributes to muscle atrophy and wasting [41–43, 73]. In this study, we identified the IL6 trans-signalling pathway as a therapeutic target for muscle-related pathologies, and specifically for DMD. Analysis of IL6ST and IL6R levels, STAT3 phosphorylation status, and changes in expression of IL6-regulated genes (upon treatment of myotubes with IL6 or IL6/IL6R complexes) strongly suggest that the trans-signalling pathway is the dominant IL6 pathway in muscle cells (**Fig. 1, Supplementary Figs. 1-3**). Furthermore, although we observed increased levels of *Il6* mRNA and phosphorylation of STAT3 in all the analysed DMD mouse models, up-regulation of the IL6 trans-signalling pathway was limited to *mdx*/IL6 mice only (**Fig. 2, Supplementary Fig. 4**). These findings could be explained by the elevated levels of systemic IL6 induced by the overexpression of human IL6 transgene in *mdx*/IL6 mice [49]. Indeed, it is plausible that the persistence of elevated levels of circulating IL6, which have been previously associated with the exacerbation of the dystrophic muscle phenotype in *mdx* mice and human patients [48, 49], can stimulate the systemic and/or local release of sIL6R, thereby promoting pathogenic IL6 action in a prodegenerative, feed-forward manner.

sIL6R is an essential mediator of the IL6 trans-signalling pathway that is primarily generated by one of two mechanisms; (i) limited proteolysis mediated by metalloproteases (such as ADAM10 and ADAM17), or (ii) alternative splicing. In the present study, we were unable to detect *Il6r* alternative splicing products in diaphragm muscle lysates from any of the various DMD mouse models tested (**Supplementary Fig. 5**). Moreover, gene expression levels of metalloproteases *(Adam10* and *Adam17),* and consequently sIL6R protein levels, were only found to be increased in *mdx*/IL6 mice (**Fig. 2**). Additionally, *Il6r* was found to be expressed at very low levels in satellite cells, when compared to *Il6st* (**Supplementary Fig. 3**). Taken together, we speculate that these observations might explain some of the differences in severity between the mildly affected *mdx* mice and the more severe pathology in *mdx*/IL6 mice (that more closely recapitulate the pathogenic features of DMD patients). It was previously shown that high levels of IL6, with chronic activation of STAT3, can lead to the exhaustion of the satellite cell pool [44, 45]. According to our results, the lack of increased levels of sIL6R in *mdx* mice, together with the fact that satellite cells do not express high levels of IL6R, might result in preservation of the satellite cells pool as STAT3 remains in an inactive state in this model. Conversely, forced expression of IL6 and STAT3 activation in *mdx*/IL6 mice leads to satellite cells exhaustion and an exacerbation of the dystrophic phenotype [49]. This is possibly due to the elevated levels of metalloproteases observed in these mice which cleave the IL6R from the cell surface and liberate sIL6R. However, in order to understand if the differences in severity between humans and *mdx* mice are related to the lack of IL6 trans-signalling pathway up-regulation in the dystrophic mouse model, it will be important to test the levels of sIL6R in DMD patient samples in future studies.

Several therapeutic strategies have been developed to counteract the deleterious effects of cytokines such as IL6, most notably the use of monoclonal antibodies targeting IL6, IL6R, or IL6/IL6R complexes [74]. However, monoclonal antibodies that target IL6 or IL6R cannot distinguish between the classical and trans-signalling pathways, resulting in blockade of both, and therefore compromise the antiinflammatory properties of the IL6. Moreover, the use of these biologics is subject to other limitations such as an increased risk of infections and malignancy [75]. Alternatively, decoy receptors (i.e. soluble cytokine receptors lacking their transmembrane and signalling domains) have also been used to block cytokines [76] although these typically exhibit very short serum half-lives (if they are not conjugated to an Fc domain), which limits their therapeutic utility [77].

The technology presented here (IL6ST-EVs) is designed to specifically block the IL6 trans-signalling pathway, without affecting the anti-inflammatory properties of the cytokine (i.e. mediated by classical signalling), as IL6ST is only able to bind IL6 when complexed with sIL6R [26]. Furthermore, EVs are expected to provide three additional advantages over the aforementioned strategies; (i) improved anti-inflammatory and pro-regenerative activities (if using EVs derived from cell types such as mesenchymal stem cells) [7, 9], (ii) the ability to cross biological barriers and be targeted to disease sites, and (iii) EVs can serve as a modular platform that enables combinatorial therapy by targeting multiple pathogenic mechanisms (e.g. either by expressing multiple decoy receptors, co-delivering therapeutic oligonucleotides or small molecule drugs), together with simultaneous co-expression of targeting ligands (to enhance tissue targeting).

Of note, endogenous EVs were shown to act as decoys for cancer treatments in some instances. One of these examples, CD20+ EVs from B cell lymphoma were found to block binding of anti-CD20 to B cells [78]. Similarly, it was found that expression of HER2 (human epidermal growth factor receptor 2) on EVs from breast cancer cells, acted as a decoy for anti-HER2 therapy [79]. Furthermore, two recent publications identified the production of endogenous decoy extracellular vesicles as a protective mechanism in response to complement-mediated cytotoxicity or bacterial toxins [80, 81]. This reveals a natural role for decoy extracellular vesicles in live organisms, including patients, and demonstrates the therapeutic potential of using these natural carriers as delivery vehicles for decoy receptors. The utility of engineered decoy receptor EVs has been further explored in a parallel study from our groups (manuscript in submission), were we were able to demonstrate improved disease phenotype in three inflammatory mouse models (systemic-, intestinal-, and neuro-inflammation).

In this study, EVs were isolated by UF-SEC-LC, a method readily scalable for the generation of large quantities of engineered, therapeutic EVs [55]. The activity of IL6ST-EVs was optimized through the addition of a GCN4 dimerization domain and the N-terminal domain of syntenin-1, which resulted in improved loading of EVs with decoy receptors and enhanced activity in reporter cell lines (**Fig. 3**). Importantly, the higher potency achieved with 2^nd^ generation IL6ST-EVs, may enable the administration of lower doses of IL6ST-EVs, thereby minimizing the potential toxicity resulting from treatments with this strategy, and reducing cost.

Experiments performed in myocytes, demonstrated that 2^nd^ generation IL6ST-EVs were able to effectively decrease the STAT3 phosphorylation in both C2C12 myoblasts and myotubes (**Fig. 4A-B**). Moreover, it has previously been shown that IL6 impairs myogenic differentiation [82], which we also observed upon treating C2C12s with IL6/IL6R complexes (**Fig. 4C**). Importantly, treatment with IL6ST-EVs was able to partially reverse this blockade of myogenic differentiation induced by IL6/IL6R complexes (**Fig. 4C-D**).

Here, we have demonstrated successful delivery of IL6ST-EVs to multiple muscle tissues upon subcutaneous administration. Nevertheless, unwanted accumulation of nanoparticles (such as EVs) in non-target organs (specifically lungs, liver, and spleen) remains a major challenge when targeting difficult to reach tissues such as muscle and brain in other contexts, as this serves to limit the effective dose that reaches these target tissues (**Supplementary Fig. 7**). In this respect, co-expressing a muscle targeting ligand (such as M12 peptide) on the EV surface, might improve accumulation in muscle [83, 84], and hopefully overcome the limitation of poor accumulation in diaphragm that we observed.

*In vivo* activity of IL6ST-EVs was confirmed by detecting a decrease in the STAT3 phosphorylation in quadriceps and gastrocnemius muscles of *mdx*/IL6 mice, two of the tissues where IL6ST-EVs accumulated (**Fig. 5**). We speculate that transient inhibition of STAT3, might contribute to a preservation of the muscle satellite cell pool and prevent dystrophy-associated stem cell depletion [44, 45]. Future studies should address the effect of transient inhibition of STAT3 with IL6ST-EVs, on satellite cell function.

Multiple studies have identified a role for the IL6 cytokine in the progression of DMD [44, 45, 47–50], but experiments that have demonstrated positive therapeutic effects of targeting this pathway utilized monoclonal antibodies against the IL6R, which do not discriminate between the IL6 signalling pathways [48, 51]. To our knowledge, this is the first study to demonstrate the importance of the IL6 trans-signalling pathway in muscle-related pathologies. Additionally, our findings show that IL6ST-EVs, might be beneficial in counteracting the pathological effects of the IL6 trans-signalling pathway in diseases such as DMD, while not interfering with the classical signalling properties of this cytokine. We hope that the engineered EV technology described here might serve as a decoy receptor platform amenable to the targeting of other molecules beyond IL6.

## Acknowledgements

This work was supported by grants from Oxford University Press John Fell Fund (awarded to TCR and IM), The University Challenge Seed Fund (UCSF) and MRC Confidence in Concept (to MJAW and SELA), Evox Therapeutics (to MJAW and SELA), Muscular Dystrophy Association (to MJAW and MC), Muscular Dystrophy UK (to MJAW, MC, and IM), Ricerca Finalizzata (to AM), Ateneo Sapienza (to AM), Swedish Medical Research Council (to SELA), and Swedish Strategic Foundation of Medical Research - SSF-IRC, FormulaEx (to SELA). AG is an International Society for Advancement of Cytometry (ISAC) Marylou Ingram Scholar (2019-2023).

## Conflict of Interest Statement

MJAW and SEA are founders of, and consultants for, Evox Therapeutics. IM, AG, and DG are consultants for Evox Therapeutics. MJAW, SEA, JZN, OW, AG, and DG are shareholders in Evox Therapeutics. The remaining authors declare no conflicts of interest.

