## Supplementary information for "Engineered extracellular vesicle decoy receptor-mediated modulation of the IL6 trans-signalling pathway in muscle"

**Table S1. List of antibodies**

| **Flow cytometry antibody** | **Source** | **ID/ Manufacturer** | **Dilution** |
|---|---|---|---|
| PE anti-mouse CD130 (IL6ST) | Rat | #149403<br>Biolegend | 1:80 |
| APC anti-mouse/rat CD126 (IL6R) | Rat | #115811<br>Biolegend | 1:20 |
| PE Rat IgG2b κ Isotype control | Rat | #400607<br>Biolegend | 1:80 |
| APC Rat IgG2b κ Isotype control | Rat | #400611<br>Biolegend | 1:20 |
| **WB primary antibody** | **Source** | **ID/ Manufacturer** | **Dilution** |
| Phospho-STAT3 (Tyr705) | Rabbit monoclonal | #9145<br>Cell Signalling Technology | 1:2000<br>(chemiluminescence) |
| Total STAT3 | Rabbit monoclonal | #8232<br>Cell Signalling Technology | 1:1000<br>(fluorescence) |
| GAPDH | Mouse monoclonal | #MAB5718<br>R&D Systems | 1:5000<br>(fluorescence) |
| Vinculin (VCL) | Mouse monoclonal | #V9131<br>Sigma Aldrich | 1:100,000<br>(fluorescence) |
| IL6Rα | Mouse monoclonal | #sc-374259<br>Santa Cruz Biotechnology | 1:1000<br>(chemiluminescence) |
| gp130 (IL6ST) | Goat polyclonal | #AF468<br>R&D Systems | 1:2000<br>(fluorescence) |
| Calnexin | Rabbit polyclonal | #ab22595<br>Abcam | 1:1000<br>(chemiluminescence) |
| Alix | Mouse monoclonal | #ab117600<br>Abcam | 1:1000<br>(chemiluminescence) |
| Syntenin-1 | Mouse monoclonal | #TA504796<br>OriGene | 1:1000<br>(chemiluminescence) |
| CD81 | Rabbit monoclonal | #ab109201<br>Abcam | 1:1000<br>(chemiluminescence) |
| **WB Secondary antibody** | **Source** | **ID/ Manufacturer** | **Dilution** |
| Anti-mouse IgG, HRP-linked | Horse | #7076<br>Cell Signalling Technology | 1:3000 |
| Anti-rabbit IgG, HRP-linked | Goat | #7077<br>Cell Signalling Technology | 1:3000 |
| Anti-goat, DyLight 800 | Rabbit | #072-07-13-06<br>KPL | 1:5000 |
| Anti-mouse, IRDye 800 CW | Goat | #926-32210<br>LI-COR | 1:5000 |
| Anti-rabbit, IRDye 800 CW | Donkey | #926-32213<br>LI-COR | 1:5000 |
| **Immunofluorescence antibody** | **Source** | **ID/ Manufacturer** | **Dilution** |
| Anti-mouse MHC | Mouse monoclonal | #MF 20<br>DSHB | 1:20 |
| Anti-mouse IgG Alexa 488 | Goat polyclonal | #A10680<br>Life Technologies | 1:500 |

**Table S2. List of primers**

| Mouse Primers | Forward sequence (5'-3') | Reverse sequence (5'-3') |
|---|---|---|
| ***Il6*** | TAGTCCTTCCTACCCCAATTTCC | TTGGTCCTTAGCCACTCCTTC |
| ***Socs3*** | CACAGCAAGTTTCCCGCCGCC | GTGCACCAGCTTGAGTACACA |
| ***Il6r*** | TGTCAACGCCATCTGTGAGTGG | ACTTTCGTACTGATCCTCGTGG |
| ***Il6st*** | CTGCTAGACTCGGAGGAGC | GCTGACTGCAGTTCTGCTTGA |
| ***Adam10*** | ATGGTGTTGCCGACAGTGTTA | GTTTGGCACGCTGGTGTTTTT |
| ***Adam17*** | AGGACGTAATTGAGCGATTTTGG | TGTTATCTGCCAGAAACTTCCC |
| ***Hprt*** | TCAGTCAACGGGGGACATAAA | GGGGCTGTACTGCTTAACCAG |
| **Splicing of IL6R** | | |
| **Mouse exon 4** (alternative splicing) | ACAGTGTGGGAAGCAAGTCC | - |
| **Mouse exon 6** (alternative splicing) | AAGGAGGAGCTTGACCTTGG | - |
| **Mouse exon 10** (alternative splicing) | - | GTGGAGGAGAGGTCGTCTTG |
| **Human exon 5** (alternative splicing) | CTCAGTGTCACCTGGCAAGA | - |
| **Human exon 10** (alternative splicing) | - | GTGGGGAGATGAGAGGAACA |

**Table S3. List of RT-qPCR primers and gBlock templates for absolute quantification**

| gBlock | Gene Fragment (5'-3') and primers |
|---|---|
| **Mouse *Il6r*** | CTTCATCATCCTGAGACTCAAGCAGAAATGGAAGTCAGAGGCTGAGAAGGAAAGCAAGACGACCT<br>CTCCTCCACCCCCACCGTATTCCTTGGGCCCACTGAAGCCGACCTTCCTTCTGGTTCCTCTC<br>Forward primer (5'-3'): CCTGAGACTCAAGCAGAAATGG<br>Reverse primer (5'-3'): AGAAGGAAGGTCGGCTTCAGT |
| **Mouse *Il6st*** | GCGTGCAGCTCACCTGCAACATCCTGTCCTTTGGGCAGATCGAGCAGAATGTGTATGGAGTCACCA<br>TGCTTTCAGGCTTTCCTCCAGATAAACCTACAAATTTGACTTGCATTGTGAATGAGGGGAAGAATAT<br>Forward primer (5'-3'): CTTTGGGCAGATCGAGCAGAA<br>Reverse primer (5'-3'): CCCTCATTCACAATGCAAGTCA |
| **Mouse *Hprt*** | ACTGAAGAGCTACTGTAATGATCAGTCAACGGGGGACATAAAAGTTATTGGTGGAGATGATCTCT<br>CAACTTTAACTGGAAAGAATGTCTTGATTGTTGAAGATATAATTGACACTGGTAAAACAATGCAAA<br>CTTTGCTTTCCCTGGTTAAGCAGTACAGCCCCAAAATGGTTAAGGTTGCAA<br>GCTTGCTGGTGAAAAGGACCTCTCGAAGTGTTGGATACAGGCCAGACT<br>Forward primer (5'-3'): TCAGTCAACGGGGGACATAAA<br>Reverse primer (5'-3'): GGGGCTGTACTGCTTAACCAG |

**Supplementary Figures**

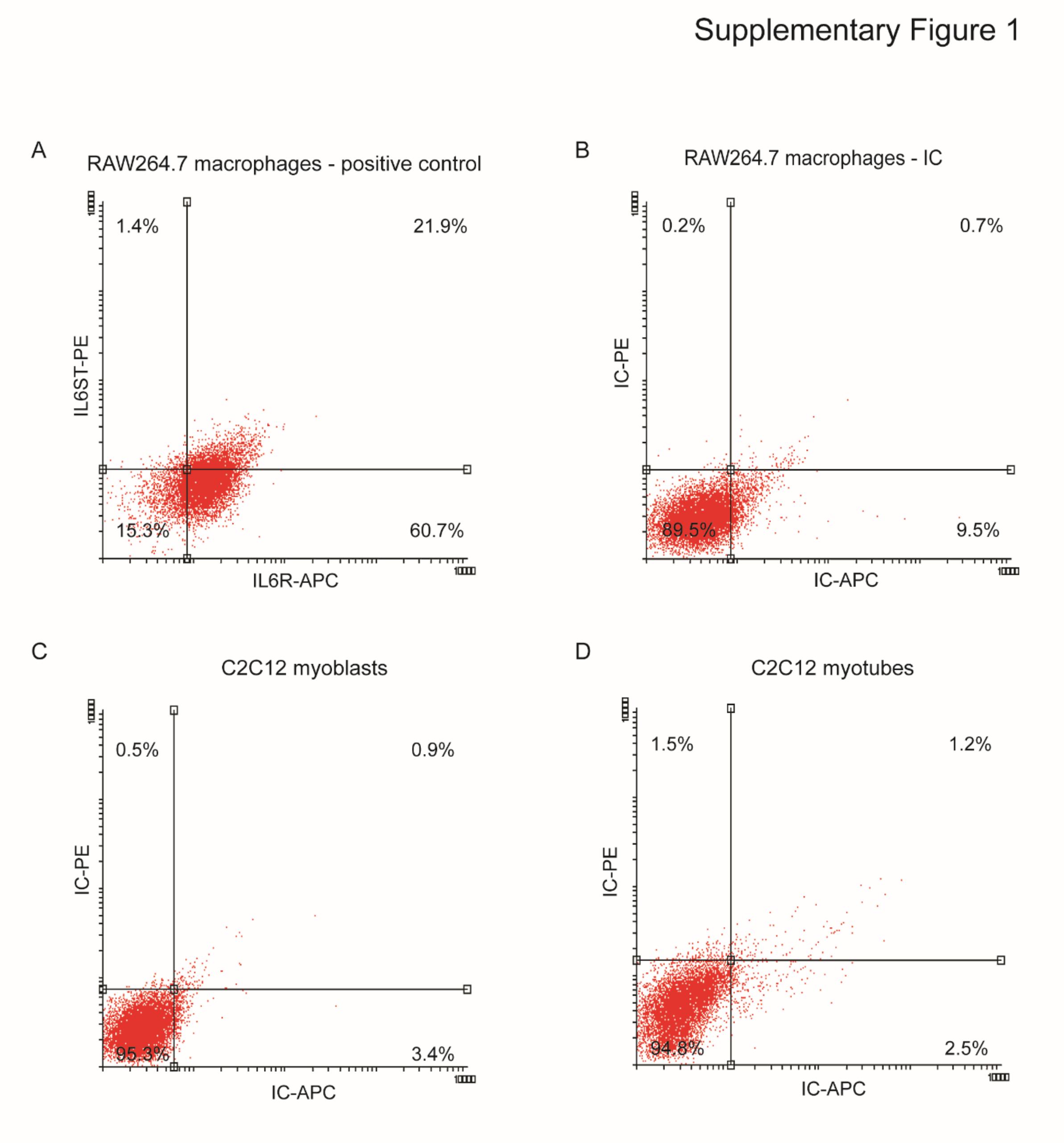

**Supplementary Figure 1. Quality control analysis for flow cytometry analysis of IL6R and IL6ST expression.**

RAW264.7 macrophages were incubated with anti-IL6R APC and anti-IL6ST PE, and the expression of IL6R and IL6ST was assessed by flow cytometry. **(A)** Two-parameter density plot for IL6R and IL6ST expression in RAW264.7 macrophages. **(B)** Two-parameter density plot obtained with incubation of RAW264.7 macrophages with the corresponding isotype control (IC) antibodies. **(C-D)** Two-parameter density plot obtained with incubation of C2C12 myoblasts **(C)** and myotubes **(D)** with the corresponding isotype control (IC) antibodies. Gating for the two-parameter density plots was set based on the background staining of the isotype controls.

Supplementary Figure 2

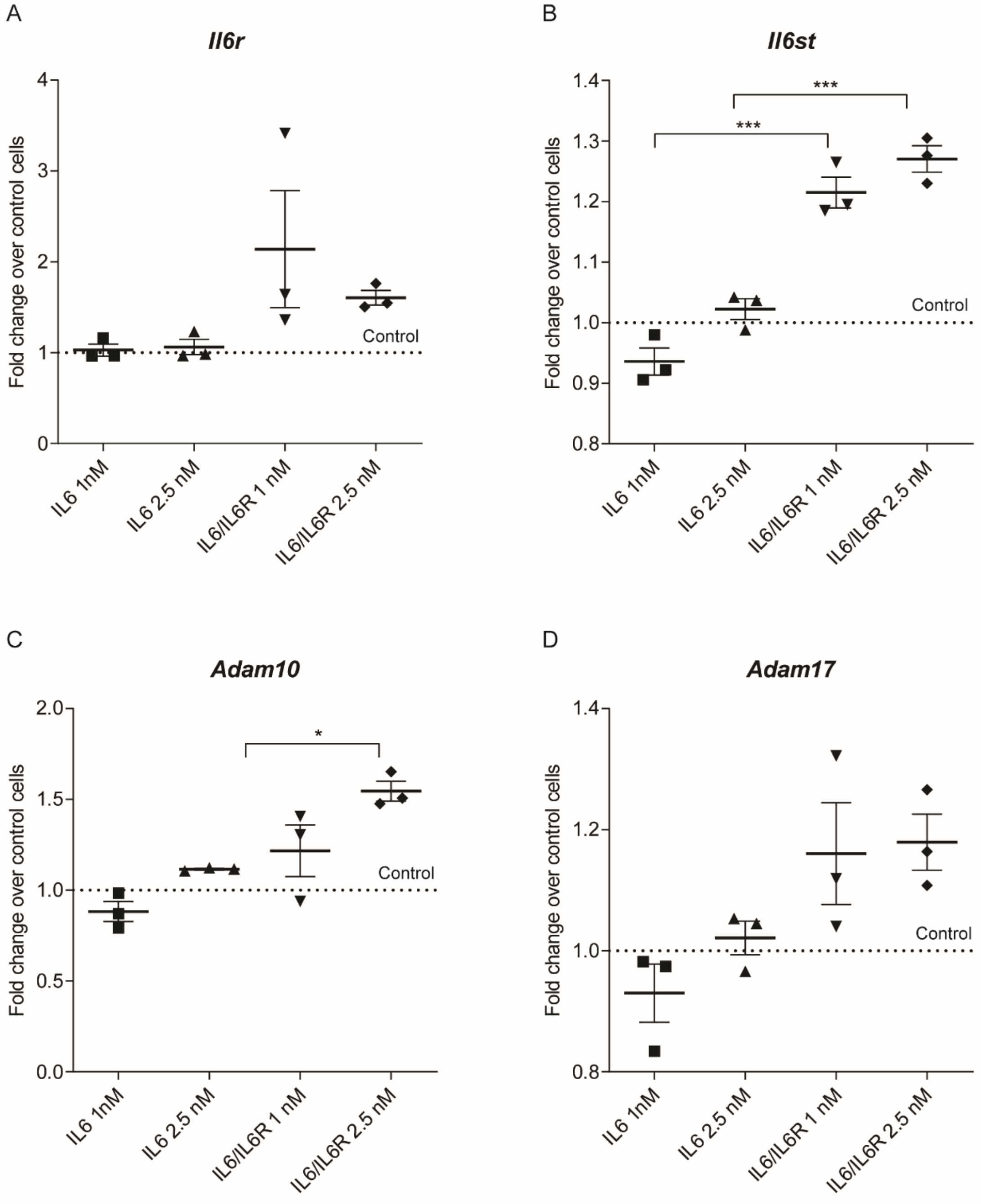

**Supplementary Figure 2. Activation of the IL6 classical and trans-signalling pathways in C2C12 myotubes.**

C2C12 myotubes were treated for 24 hours with recombinant IL6 or IL6/IL6R complexes (1 or 2.5 nM, as indicated) and mRNA expression determined by RT-qPCR for **(A)** *Il6r*, **(B)** *Il6st*, **(C)** *Adam10,* and **(D)** *Adam17*. The dotted lines indicate the levels of expression in untreated control cells. Gene-of-interest expression was normalized to the levels of a stable reference gene; *Hprt*. Values are mean±SEM, $n$=3 independent experiments, *$P$<0.05, ***$P$<0.001 (one-way ANOVA with Bonferroni *post hoc* test).

Supplementary Figure 3

A

| Transcript | Total number of copies | |
|---|---|---|
| | **C57/BL10** | ***mdx*** |
| ***Il6r*** | $3.93x10^5$ | $8.49x10^5$ |
| ***Il6st*** | $8.97x10^6$ | $2.52x10^7$ |
| ***Hprt*** | $4.92x10^6$ | $1.1x10^7$ |

B

| Transcript | Fold change over *Hprt* | |
|---|---|---|
| | **C57/BL10** | ***mdx*** |
| ***Il6r*** | 0.0799 | 0.0772 |
| ***Il6st*** | 1.82<br>(↑23.6x relative to IL-6R) | 2.29<br>(↑29.7x relative to IL-6R) |

**Supplementary Figure 3. Absolute quantification of *Il6r* and *Il6st* transcript levels in muscle satellite cells.**

**(A)** Absolute quantification of *Il6r, Il6st,* and *Hprt* levels in muscle satellite cells from WT (C57/BL10) or *mdx* mice. **(B)** Relative levels of *Il6r* and *Il6st* transcripts normalized to the expression of the *Hprt* reference gene. *Il6st* is expressed at 23.6-29.7 fold higher levels than *Il6r* in satellite cells.

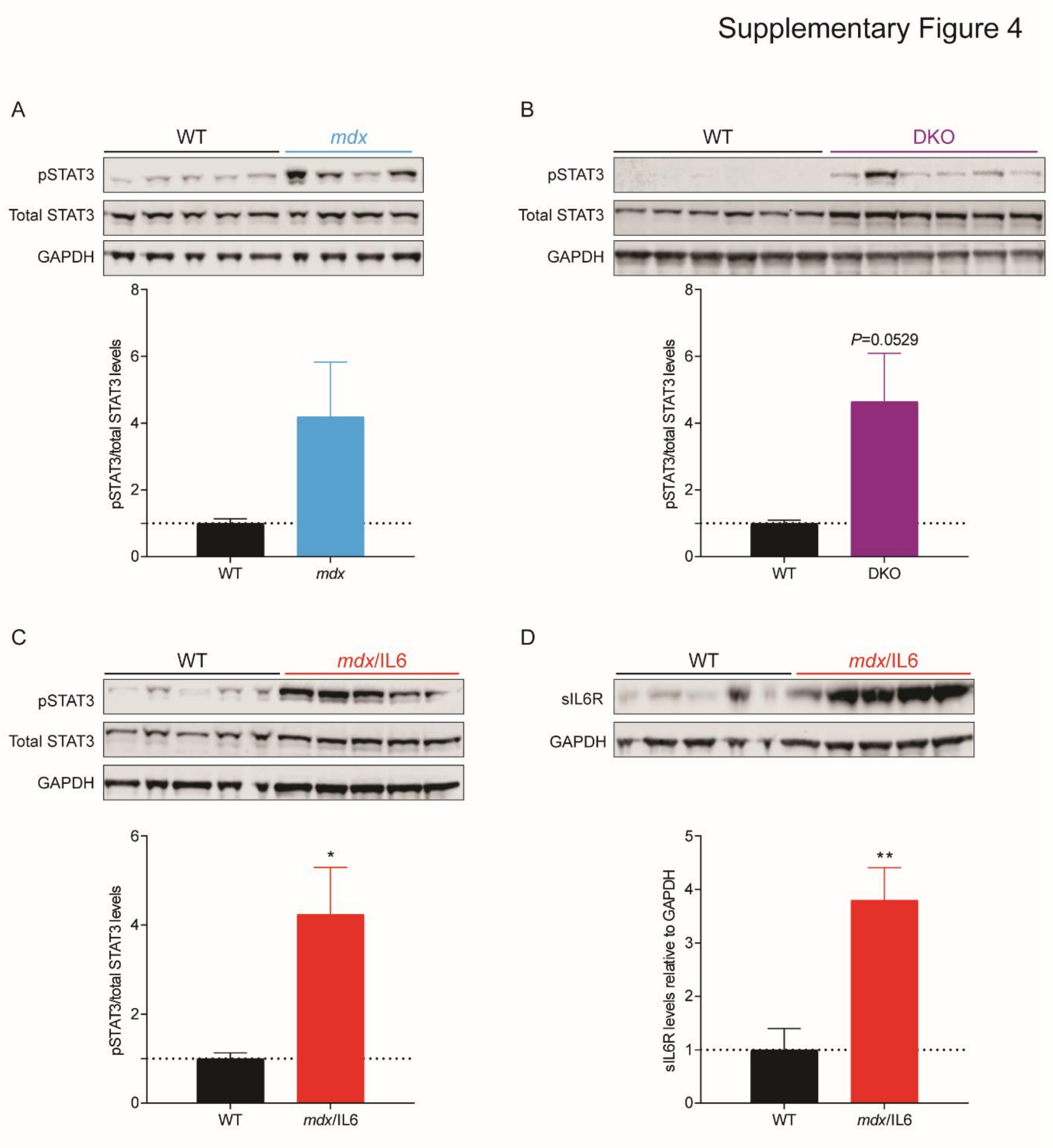
Supplementary Figure 4
A
WT
mdx
pSTAT3
Total STAT3
GAPDH
8
6
4
2
0
pSTAT3/total STAT3 levels
WT
mdx
B
WT
DKO
pSTAT3
Total STAT3
GAPDH
P=0.0529
8
6
4
2
0
pSTAT3/total STAT3 levels
WT
DKO
C
WT
mdx/IL6
pSTAT3
Total STAT3
GAPDH
*
6
4
2
0
pSTAT3/total STAT3 levels
WT
mdx/IL6
D
WT
mdx/IL6
sIL6R
GAPDH
**
5
4
3
2
1
0
sIL6R levels relative to GAPDH
WT
mdx/IL6

**Supplementary Figure 4. The IL6 pathway is altered in hind limb muscles of DMD mouse models.** Hind limb muscles of *mdx*, DKO, and *mdx*/IL6 mice were analysed at 6.5 weeks of age for *mdx* and DKO mice, and at 12 weeks of age for *mdx*/IL6 mice. STAT3 activation in the **(A)** quadriceps of *mdx*, **(B)** tibialis anterior (TA) of DKO**,** and **(C)** quadriceps of *mdx*/IL6 mice, as assessed by western blot (WB) for active phospho-STAT3 (pSTAT3, Tyr705), total STAT3, and GAPDH. **(D)** sIL6R levels in the quadriceps of *mdx*/IL6 mice, assessed by WB. Quantification of WB signal from multiple independent experiments is shown for each analysis. Values are mean+SEM, $n$=4-6 independent experiments for WB, *$P$<0.05, **$P$<0.01 (unpaired Student's $t$-test).

Supplementary Figure 5

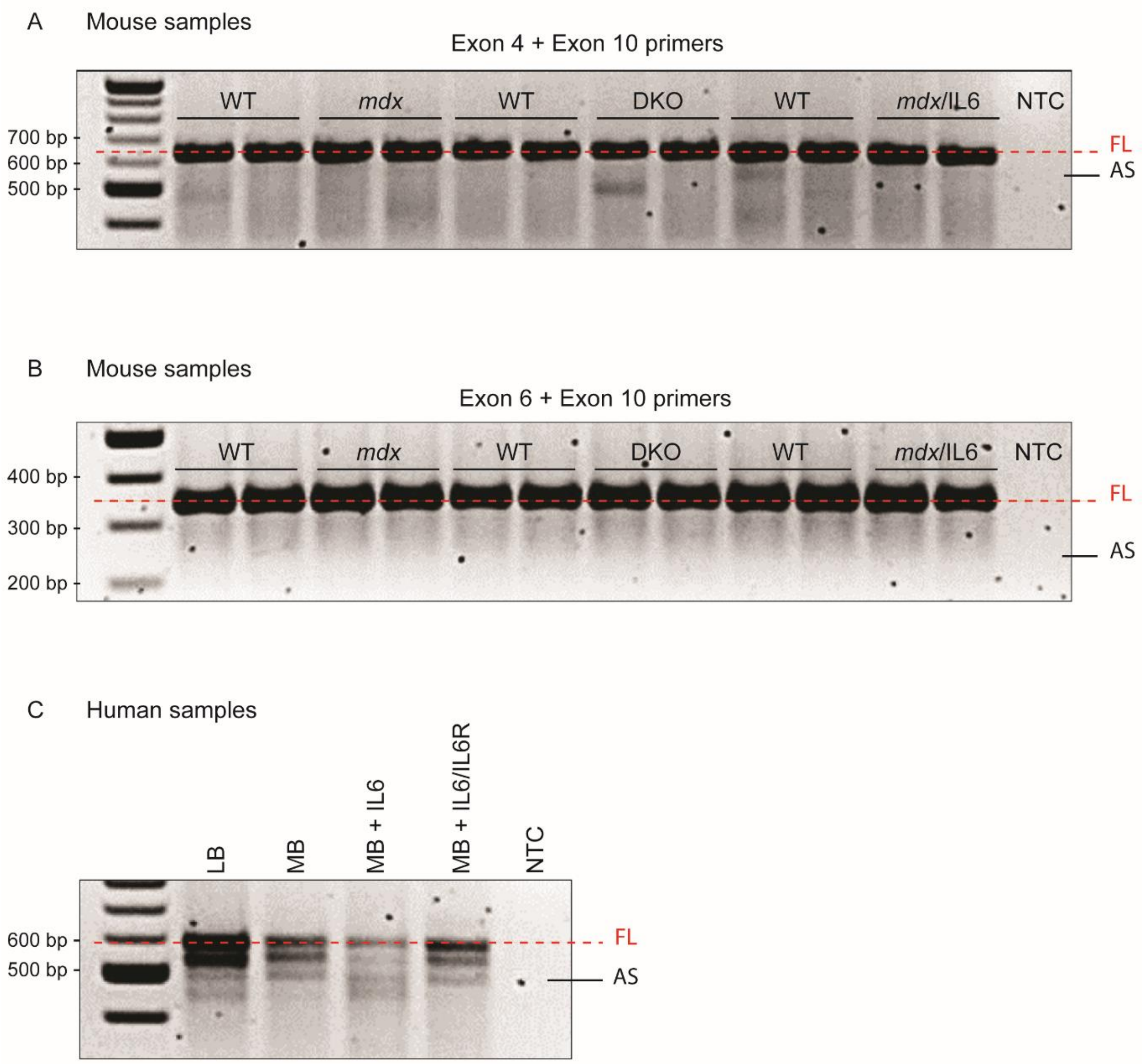

**Supplementary Figure 5. *Il6r* alternative splicing is not detected in mouse models of DMD.**

The mouse *Il6r* gene contains 10 exons, with exon 9 encoding the membrane-spanning domain. RT-PCR with primers which span this region **(A)** exon 4 to exon 10, and **(B)** exon 6 to exon 10, was performed to assess alternative splicing patterns in diaphragm lysates from WT, *mdx*, DKO, and *mdx*/IL6 mice. **(C)** RT-PCR with primers which flank the human IL6R transmembrane domain was performed as a positive control for alternative splicing. The expected amplicon lengths are indicated for full length (FL) and alternatively spliced (AS) products. No-template control (NTC) samples served as negative controls.

Supplementary Figure 6

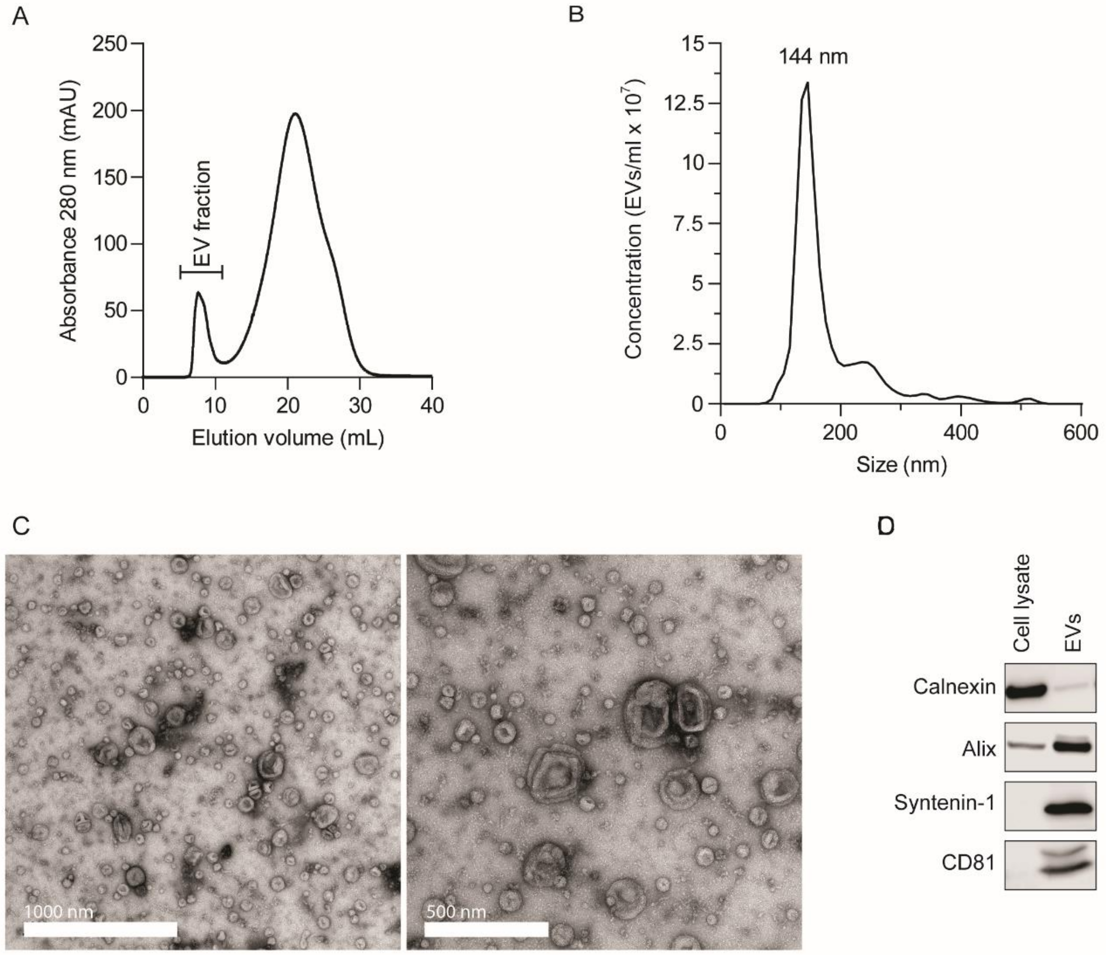

**Supplementary Figure 6. EV isolation and characterization.**

**(A)** EVs were isolated by UF-SEC-LC, and monitored by UV spectroscopy (absorbance at 280 nm). The fractions containing EVs are indicated. **(B)** Size distribution plot of EVs from HEK293T cells, as determined by nanoparticle tracking analysis (NTA). The graph is representative of *n*=3 independent experiments. **(C)** Transmission electron microscopy (EM) images of HEK293T EVs. **(D)** Western blot for calnexin, alix, syntenin-1 and CD81 performed on HEK293T cell lysate and isolated EVs.

Supplementary Figure 7

A

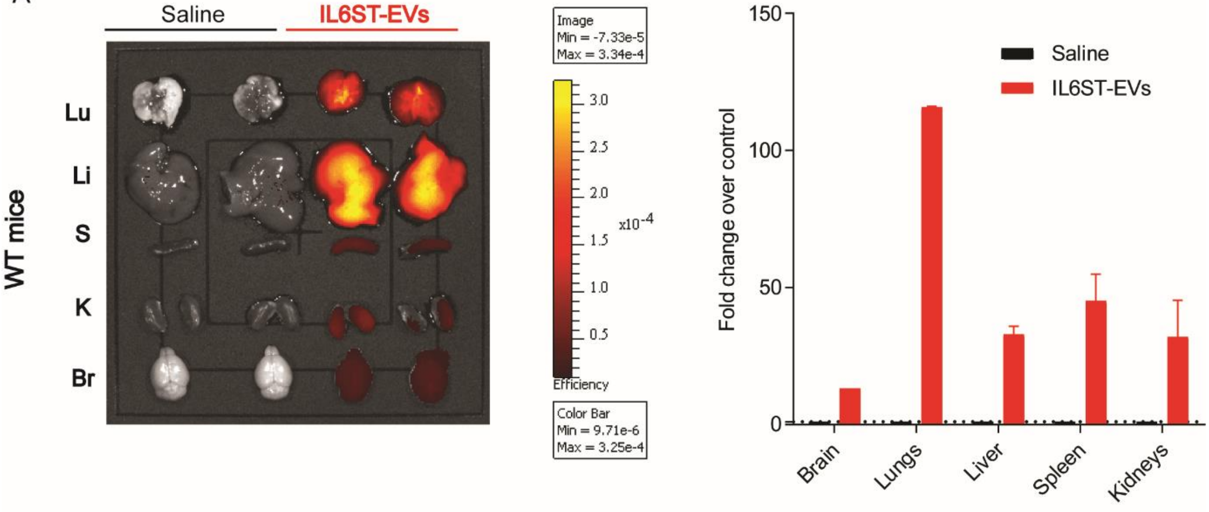

B

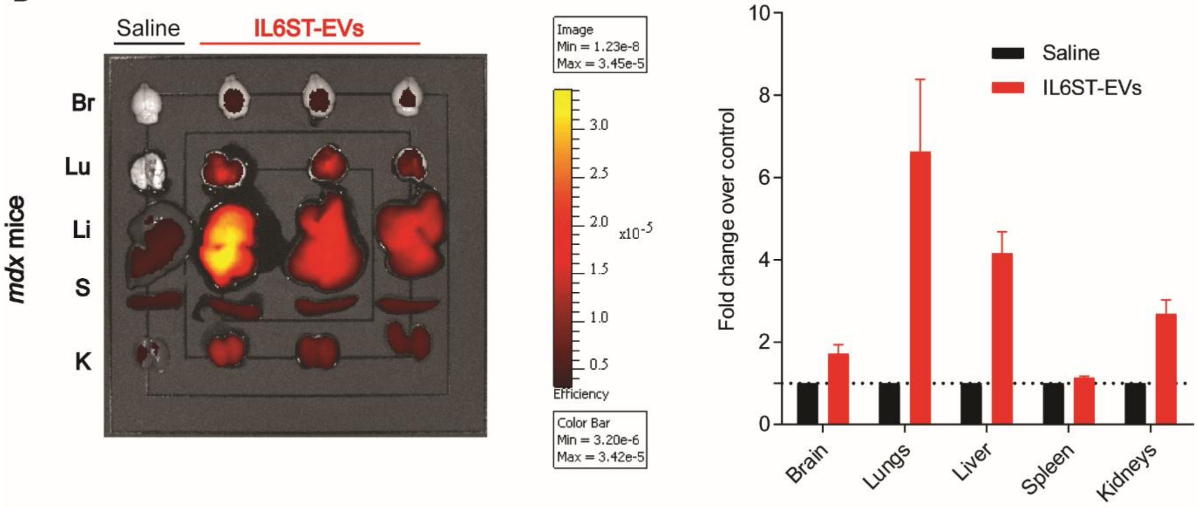

**Supplementary Figure 7. IL6ST-EVs biodistribution in peripheral organs.**

**(A)** WT or **(B)** *mdx* mice were subcutaneously injected with DiR-labelled MSC $2^{nd}$ generation IL6ST-EVs ($8x10^9$ EVs/g) and images were acquired 24 hours after injection using an IVIS Lumina imager. Each image includes lungs (Lu), liver (Li), spleen (S), kidneys (K) or brain (Br) from mice injected with either DiR-labelled IL6ST-EVs or saline-injected controls. Values are presented as mean+SEM of *n*=1-3 for biodistribution (as shown).

Supplementary Fig. 8

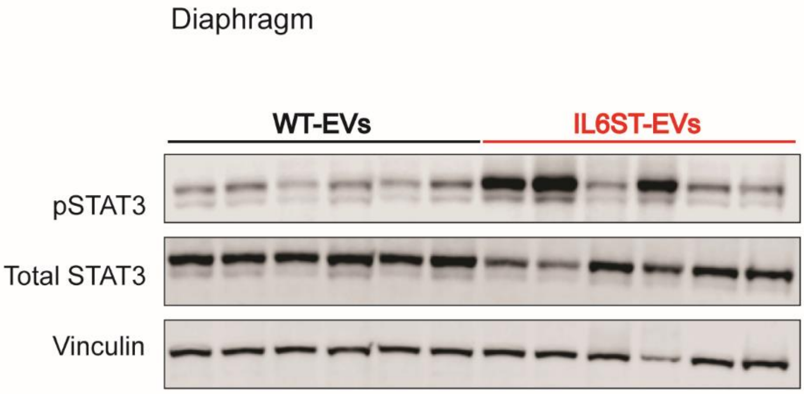

**Supplementary Figure 8. IL6ST-EVs do not decrease activation of STAT3 in the diaphragm of *mdx*/IL6 mice.**

4 weeks-old *mdx*/IL6 mice were treated with IL6ST-EVs or WT-EVs (1x$10^{10}$ EVs/g) for 2 weeks, 2 injections per week. Mice were sacrificed 3 days after the 4$^{th}$ injection, diaphragm muscles harvested, and the levels of pSTAT3, total STAT3, and vinculin determined by WB.
